# Mechanistic insights into RNA surveillance by the canonical poly(A) polymerase Pla1 of the MTREC complex

**DOI:** 10.1101/2022.07.20.500385

**Authors:** Komal Soni, Anusree Sivadas, Attila Horváth, Nikolay Dobrev, Rippei Hayashi, Leo Kiss, Bernd Simon, Klemens Wild, Irmgard Sinning, Tamás Fischer

**Affiliations:** Heidelberg University Biochemistry Center (BZH), INF 328, D-69120 Heidelberg, Germany; The John Curtin School of Medical Research, The Australian National University, Canberra, ACT 2601, Australia; European Molecular Biology Laboratory (EMBL), Meyerhofstr, 1, D-69117 Heidelberg, Germany

## Abstract

The *S. pombe* orthologue of the human PAXT complex, Mtl1-Red1 Core (MTREC), is an eleven-subunit complex which targets cryptic unstable transcripts (CUTs) to the nuclear RNA exosome for degradation. It encompasses the canonical poly(A) polymerase Pla1, responsible for polyadenylation of nascent RNA transcripts as part of the cleavage and polyadenylation factor (CPF/CPSF). In this study we identified and characterised the interaction between Pla1 and the MTREC complex core component Red1 and analysed the functional relevance of this interaction *in vivo*. Our crystal structure of the Pla1-Red1 complex showed that a 58-residue fragment in Red1 binds to the RNA recognition motif domain of Pla1 and tethers it to the MTREC complex. Structure-based Pla1-Red1 interaction mutations showed that Pla1, as part of MTREC complex, hyper-adenylates CUTs for their efficient degradation. Interestingly, the Red1-Pla1 interaction was also required for the efficient assembly of the fission yeast facultative heterochromatic islands. Together, our data suggest a complex interplay between the RNA surveillance and 3’-end processing machineries.

## Introduction

Nuclear RNA surveillance is carried out primarily by the exosome which is involved in 3’-5’ RNA degradation and maturation processes^1–3^. While export-competent mature RNAs are packaged into mRNP particles, transcripts that fail to mature properly are rapidly degraded by the nuclear RNA surveillance machinery. In addition, pervasive transcription, occurring widely in eukaryotes largely due to bidirectional promoters, produces tremendous amounts of non-coding RNA transcripts that are degraded by the RNA surveillance machinery^4–7^. These pervasive transcripts, also known as cryptic unstable transcripts (CUTs) in yeast, or promoter upstream transcripts (PROMPTs) and upstream antisense RNAs (uaRNAs) in humans, are barely detectable under steady-state conditions due to their rapid degradation^8, 9^. The herculean task of specific recognition, targeting and degradation of the wide variety of defective RNAs by the exosome is done by its so-called adaptor complexes^1, 10^. The nuclear exosome relies on the helicase activity of the DExH-box RNA helicase Mtr4 to thread single-stranded RNAs into its core, thereby making it a *bona fide* member of all adaptor complexes^11–16^. The best characterised adaptor complex is the TRAMP complex^17^, which was identified as the major exosome targeting factor for CUTs in *Saccharomyces cerevisiae*^18^.

Polyadenylation is a post-transcriptional modification of RNA involving the addition of poly(A) tails to the 3’-ends of RNA substrates^19, 20^. Following the discovery of the yeast TRAMP complex, the connection between polyadenylation-mediated destabilisation of RNAs as a signal for exosome degradation in eukaryotes was established^21, 22^. In humans, two recently identified adaptors known as the nuclear exosome targeting (NEXT)^23^ and the poly(A) tail exosome targeting (PAXT) complex^24^ are responsible for targeting early, unprocessed RNAs, PROMPTs and enhancer RNAs or transcripts with extensive poly(A) tails, respectively.

In *Schizosaccharomyces pombe*, a highly conserved MTREC (Mtl1-Red1 core)^25, 26^ or NURS (nuclear RNA silencing) complex^27^ is responsible for degradation of CUTs and meiotic mRNAs. Notably, the MTREC complex was identified in studies of the Mtr4-like protein 1 (Mtl1) and the zinc-finger protein Red1, which in turn contributed to the definition of the PAXT complex in humans^28^. MTREC is a multi-subunit complex comprising the CAP binding complex Cbc1-Cbc2-Ars2 (CBCA), Iss10-Mmi1, Red5-Pab2-Rmn1 and the canonical poly(A) polymerase Pla1, in addition to the core Mtl1 and Red1 proteins^26^. We have recently described Red1 as the central scaffolding protein of MTREC, which directly interacts with each submodule, including Pla1, and bridges them to the helicase Mtl1 for targeting of the associated RNA cargo to the exosome^29^. Importantly, Pla1 is usually part of the highly conserved cleavage and polyadenylation factor (CPF or CPSF in mammals), where it functions to add poly (A) tails during the 3’-end processing of transcripts^30–32^. Co-purification of Pla1 with the MTREC complex and the extended poly(A) tails in meiotic mRNAs and CUTs that are necessary for their efficient degradation^33–35^ therefore appear to be correlated. Consistently, Mmi1, along with Pla1 and poly(A) binding protein Pab2 have been implicated in promoting hyperadenylation of CUTs and meiotic RNAs in fission yeast ^34, 36^. In addition, Pla1 co-localised and immunoprecipitated with Red1 in an RNA-independent fashion, suggesting a physical interaction between them^35^. Furthermore, it was shown that Red1 and Mmi1 facilitate the hyper-polyadenylation of DSR-containing mRNAs by Pla1^35^ and that this hyper-adenylation is required for their efficient degradation.

Studies of Pla1 homologues from mammals (PAP) and *S. cerevisiae* (Pap1) show that the polymerase adopts a tri-partite domain architecture^37, 38^. It consists of three globular domains, an N-terminal nucleotide binding domain (NTD), a middle domain (MD) and a C-terminal RNA Recognition Motif (RRM)^39^, where the NTD and MD form the catalytic center (**Figure 1A**). Detailed kinetic and thermodynamic analyses of Pap1 suggest that large-scale domain movements in the protein are required for substrate recognition and catalysis^40–42^ and the enzyme is stabilised in a closed conformation with extensive contacts between the NTD and RRM domain upon substrate binding^40^. Sequence alignment of Pla1 from different species shows that the NTD and MD are highly conserved between yeast and mammals (66.6 % similarity), while the RRM domain is quite diverse (19.5 % similarity) (**Supplementary Figure S1**). Interestingly, the C-terminal RRM domain lacks the consensus RNA binding residues (reviewed in ^43^) and does not interact with the RNA substrate via its canonical RNA binding ′-sheet interface, but instead binds via its opposite interface^40^. In *S. cerevisiae*, the RRM domain of Pap1 is also responsible for binding to the cleavage and polyadenylation subunit Fip1 (homologous to Iss1 in *S. pombe*)^44, 45^, tethering it to the CPF. Binding of Fip1 leads to Pap1 inhibition in yeast, whereas activation has been observed in mammals and plants^44, 46, 47^.

**Figure 1.**
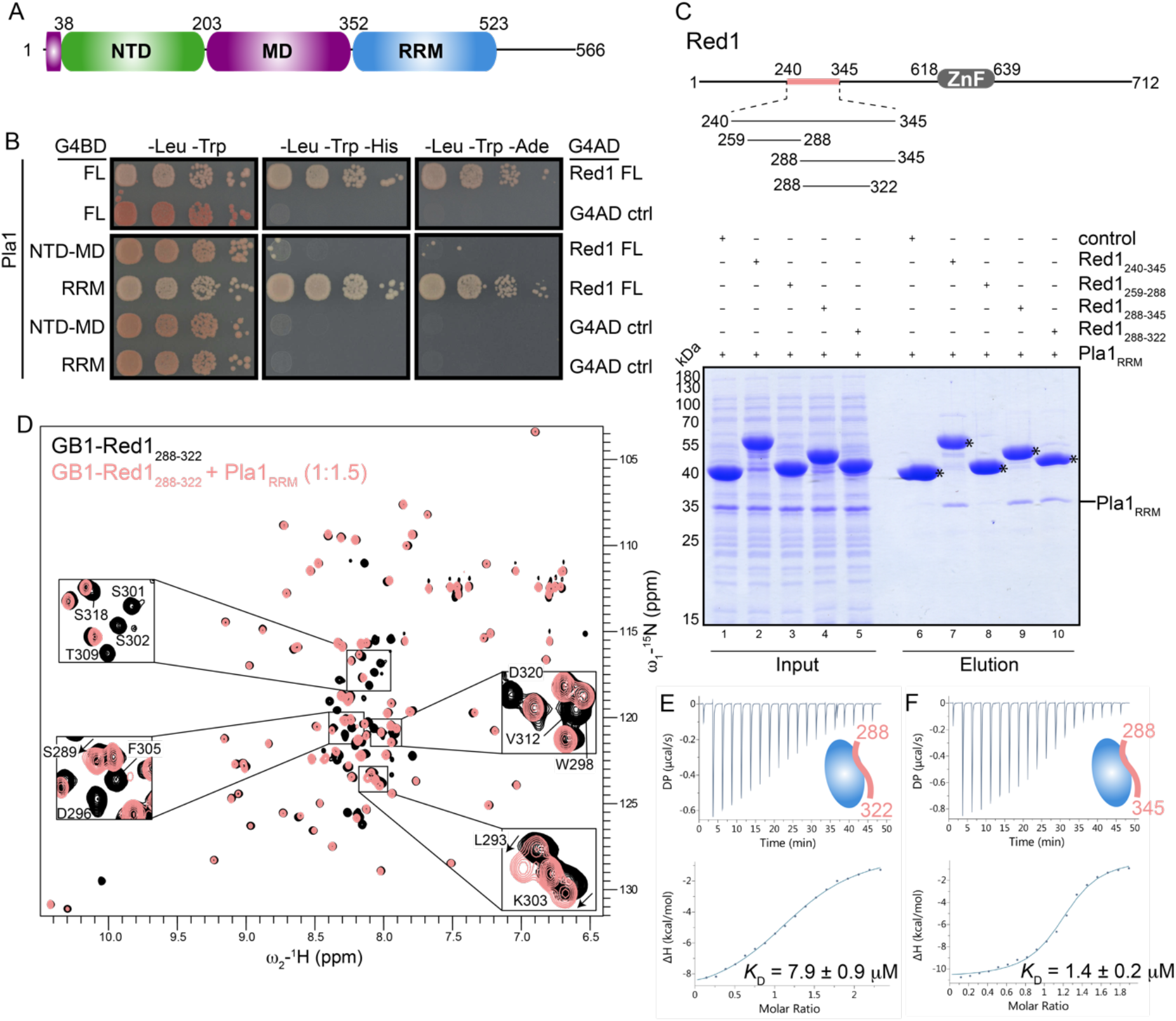
Interaction mapping between Pla1 and Red1. (A) Domain organization of Pla1, consisting of the NTD, MD and RRM domains shown in green, purple and blue, respectively. (B) Y2H experiments show that the Pla1 RRM domain is responsible for the interaction with Red1. Full-length (FL) Pla1, Pla1_NTD-MD_ or Pla1RRM constructs were fused to the Gal4 DNA binding domain (G4BD) while full-length Red1 was fused to the Gal4 activation domain (G4AD). Auto-activation controls (ctrl) for FL-Pla1 Pla1_NTD-MD_ and Pla1_RRM_ are provided. Serial dilutions of equivalent amounts of yeast were plated on double (-Leu-Trp) and triple dropout media (-Leu-Trp-His, -Leu-Trp-Ade), with growth on triple dropout media indicating an interaction between the tested proteins. (C) *In vitro* pull-down assays. Untagged Pla1_RRM_ domain was co-expressed with His_6_-MBP fusion constructs of Red1 or His_6_-MBP (control). 0.01 % of input (lanes 1-5) and 25 % of elution fractions (lanes 6-10) were separated on 12 % SDS-PAGE gel. Asterisks mark the different His_6_-MBP fusion constructs. (D) Overlay of ^1^H, ^15^N-HSQC NMR spectra of Red1_288-322_ in the absence and presence of Pla1_RRM_ domain are shown in black and salmon, respectively. Zoom-in views of Red1 residues showing chemical shift perturbations or line broadening are shown. (E and F) ITC experiments with a serial titration of Red1_288-322_ (E) or Red1_288-345_ (F) into Pla1_RRM_ domain. The calculated dissociation constants (*K*_D_) from an average of two independent measurements are shown.

While Pla1-dependent excessive polyadenylation of meiotic mRNAs that are destined for degradation^33^ and hyperadenylation of CUTs has been reported in *S. pombe*^26^ (compared to *S. cerevisiae* where MTREC is absent), the functional importance of Pla1 in MTREC has not been studied. In this study, we have used a combination of structural, biochemical and *in vivo* experiments to understand the specific role of Pla1 as an associated factor of the MTREC complex. Together with nuclear magnetic resonance spectroscopy (NMR) and isothermal titration calorimetry (ITC), we show that the C-terminal RRM domain of Pla1 specifically recognises a 58-residue fragment of Red1. We report crystal structures of Pla1 in its apo form and in complex with Red1. Specific point mutations disrupting the Pla1-Red1 interaction and deletion of the complete Pla1 interacting region in Red1 lead to the widespread accumulation of PROMPTs which also harbour shorter poly(A) tails compared to wild-type cells. Interestingly, the Pla1 “truncated” MTREC complex is also defective in the assembly of facultative heterochromatic islands at meiotic genes. Taken together, our data reveal the molecular details of recruitment of Pla1 by Red1 which leads to hyperadenylation of CUTs, serving as a signal for exosome-mediated target degradation. In addition, *in vitro* competition experiments between Red1 and Iss1, and additional interactions between MTREC and the CPF-associated factor Msi2 (homologous to CF IB in *S. cerevisiae*), indicate the existence of an intricate interaction network between CPF and the MTREC complex to control the processing and surveillance of nascent RNA transcripts.

## Results

### The C-terminal RRM domain of Pla1 mediates its interaction with Red1

We have previously shown that the Red1 fragment comprising residues 240-345 (Red1240-345) mediates interaction of Pla1 with the MTREC complex^29^, but the exact interaction region within Pla1 was not identified. Therefore, we set out to map the minimal interaction region of Pla1 with Red1 using yeast-two hybrid experiments (Y2H). Y2H assays were performed with constructs comprising the N-terminal-(NTD) and middle-domains (MD) of Pla1 (residues 1-352, NTD-MD) or RNA recognition motif (RRM) domain of Pla1 (residues 352-566) (**Figure 1B**). We found that the Pla1 RRM domain (Pla1_RRM_) is necessary and sufficient for interaction with Red1, consistent with previous reports that the RRM domain of the *S. cerevisiae* poly(A)-polymerase (Pap1) is responsible for mediating protein-protein interaction between Pap1 and its co-factor, Fip1^44,45^.

To further validate our Y2H results, we decided to reconstitute the Pla1-Red1 complex *in vitro*. To this end, we used bacterial cell lysates co-expressing His_6_-MBP fused to various Red1 fragments together with untagged Pla1_RRM_ and assessed the ability of the Red1 constructs to pull down Pla1_RRM_ (**Figure 1C**). We used partially overlapping, short fragments of Red1, comprising residues 259-288 (Red1_259-288_), 288-345 (Red1_288-345_), and 288-322 (Red1_288-322_) in addition to the originally identified fragment comprising residues 240-345 (Red1_240-345_). While Red1240-345, Red1_288-345_ and Red1_288-322_ co-purified the Pla1_RRM_ domain, Red1_259-288_ did not, suggesting that residues 288-322 of Red1 contain the Pla1 binding site. We further probed the Pla1-Red1 binding interface using NMR. A two dimensional ^1^H, ^15^N-HSQC spectrum of Red1_288-322_ shows that this region of Red1 is largely unstructured, as inferred from the low chemical shift dispersion in the ^1^H-dimension (**Figure 1D**). Subsequently, we performed NMR titrations of unlabeled Pla1_RRM_ into ^15^N-labeled Red12_88-322_ and monitored the chemical shifts using two dimensional ^1^H, ^15^N-HSQC spectra (**Figure 1D**). We observed that a majority of the backbone amide resonances of the Red1 peptide show severe line broadening while a handful show chemical shift perturbations (CSPs), both indicative of a clear binding event (**Supplementary Figure S2A, B**). To confirm whether Red1_288-322_ contains the complete Pla1 binding site, we performed NMR titrations of unlabeled Pla1_RRM_ into ^15^N-labeled Red1_288-345_ (**Supplementary Figure S2C**). Surprisingly, we observed additional residues in Red1_288-345_ which show CSPs compared to that of Red1_288-322_. To understand the affinity contribution of these additional residues in Red1_288-345_ and define the exact region of Red1 necessary and sufficient for binding to Pla1, we performed isothermal titration calorimetry. We found that while Red1_288-322_ binds to Pla1 with a *K*_D_ = 7.9 µM (**Figure 1E**), the addition of residues 323-345 in Red1_288-345_ leads to a ∼5.6-fold increase in binding affinity (*K*_D_ = 1.4 µM, **Figure 1F**). We therefore refer to the Red1 fragment comprising residues 288-345 as the complete Pla1 interaction site.

### Crystal structures of the canonical poly(A) polymerase Pla1 in its apo form and in complex with Red1

To first gain insights into the inter-domain arrangement and interactions between the NTD, MD and RRM domains of Pla1, we determined the crystal structure of the protein in its apo form. We expressed, purified and performed crystallization trials with the full-length (Pla1_FL_) and a C-terminal truncation (Pla1_Δ14_) of Pla1. Both constructs readily crystallized under a variety of conditions within 2-4 days. Crystals of Pla1_FL_ diffracted to substantially higher 1.9 Å resolution while Pla1_Δ14_ diffracted to 2.6 Å resolution.

The crystal structure of the full-length protein shows a tripartite domain architecture of Pla1 (**Figure 2A**). The overall topology of the three domains is largely similar to that of the Pla1 homologs (**Supplementary Figure S3A**). Briefly, the N-terminal catalytic domain is homologous to the catalytic domain of other nucleotidyl-transferases forming a five stranded mixed β-sheet along with four α-helices^48, 49^. The middle domain is formed by a four-helix bundle which is capped by the N-terminus of Pla1. The C-terminal RRM domain contains additional secondary structure elements extending the canonical βαββαβ topology of RRMs^43^. It comprises additional helices α14, α15 and β-strands β8, β9 in the extended loop L1 connecting the canonical RRM strands β7 and β10 compared to its homologs (**Supplementary Figure S3B**). In our crystal structure, 18 residues of loop L1, loop L2 connecting β10 and α16 and 25 residues at the C-terminus are disordered. Interestingly, residues 415-434, comprising helix α14 and β-strand β9, partially block the canonical RNA binding interface of the RRM domain on one side^39^, while the other side is blocked by residues 523-532 belonging to the C-terminus of the protein. The helix α15 of the RRM domain is positioned away from the β-sheet interface facing the back side of the RRM, to form an interface with the Pla1NTD (**Figure 2A** **and Supplementary Figure S4A**).

**Figure 2.**
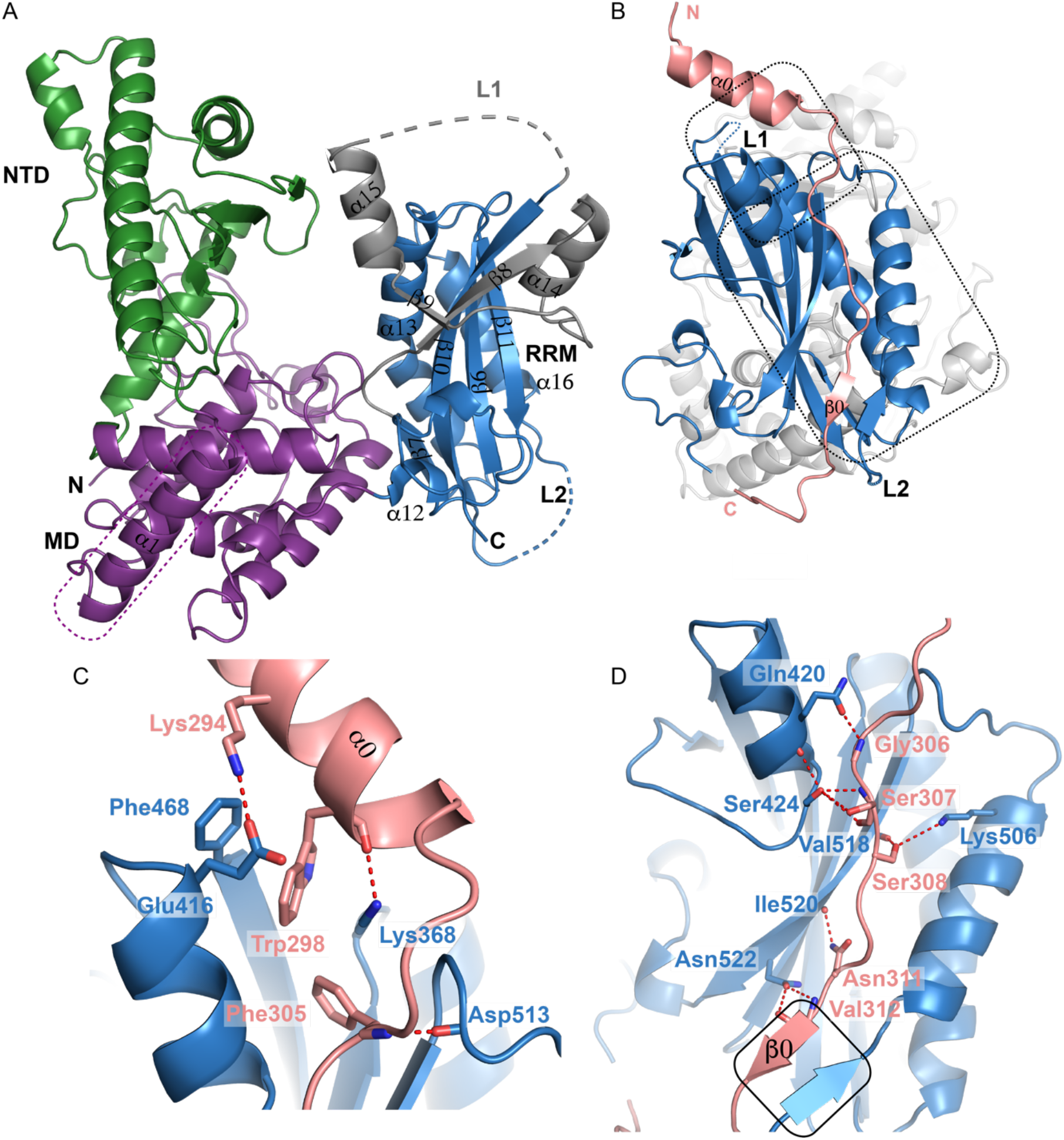
Crystal structures of Pla1 in its apo form and in complex with Red1. (A) The overall architecture of Pla1 in its apo form is shown. Color scheme for Pla1 domains are the same as Figure 1A. The N- and C-termini of the protein and loops L1 (shown in grey) and L2 of RRM domain are marked. The Pla1 N-terminal α1, which forms a part of the MD, is boxed in purple. The secondary structure elements of the RRM domain comprising helices α12-α16 and β-strands β6-β11 are marked. (B) Crystal structure of Pla1-Red1 complex. Red1 and Pla1_RRM_ are shown in salmon and blue, respectively, while the NTD and MD are colored in grey. The Red1 N- and C-termini, its secondary structure elements α0 and β0, along with the Pla1_RRM_ domain loops L1 and L2 are marked. The two Red1 interaction interfaces are boxed. (C) and (D) Zoom-in views of residues involved in the Pla1-Red1 interaction. Polar contacts are shown by red dotted lines. The β-zipper formed between Red1 β0 and Pla1 residues Arg491-Asp493 is boxed in black.

A large interface is formed by a network of interactions between the three domains of the protein burying a total surface area of ∼2982 Å^2^. The tripartite domain architecture of Pla1 is held together by hydrogen bonds between the interface residues of the domains (**Supplementary Figure S4A**). A comparison of our crystal structures of Pla1_FL_ and Pla1_Δ_14 shows that the individual domains have undergone significant movements. An alignment based on the middle domains illustrates that the N-terminal has a modest rotation of 2.95° with a 1.13 Å displacement, while the C-terminal RRM domain has a more pronounced rotation of 9.8° and 2.26 Å displacement (**Supplementary Figure S4B**). Pap1 is also known to undergo large-scale domain motions around the defined hinges connecting the NTD and MD, and the MD and RRM domains^40, 42^, which are deemed necessary for proper functioning of the enzyme.

To understand the molecular details of the Red1-Pla1 interaction, we determined the crystal structure of the complex. Since Red1 binds to Pla1_RRM_ with low micromolar affinity, we reasoned that the complex should be stably purified using size exclusion chromatography. Indeed, we could reconstitute a highly pure complex which was subjected to co-crystallization trials (**Supplementary Figure S5A**). Co-crystals of the complex appeared within three weeks and diffracted to 2.81 Å resolution. The asymmetric unit contains two molecules of the Pla1-Red1 heterodimer which exhibit small differences between them, as indicated by a root mean square deviation (RMSD) of 0.96 Å. Although the Pla1-Red1 interaction interface is conserved between the two molecules, the overall density for the second heterodimer is much weaker and therefore our structural analyses are based on the first molecule. As in the apo structure, part of loop L1, the complete loop L2 and the C-termini of Pla1_RRM_ domain are disordered. RRM domain α15, which forms an interface with the NTD in our apo structure, is also partly disordered. In addition to these, the Red1 C-terminus, spanning residues 323-345, is flexible and therefore not observed in the crystal structure of the complex. A superposition of the apo Pla1_FL_ and the Pla1-Red1 complex shows that the Pla1NTD has a rotation of 4.98° with a 1.01 Å displacement, while the Pla1_RRM_ domain has an angular rotation of 8.06° with a 1.63 Å displacement, again indicative of the flexibility between the domains (**Supplementary Figure S5B**).

**Table 1.** X-ray crystallography data collection and refinement statistics.

|  | Pla1<sub>FL</sub> | Pla1<sub>Δ14</sub> | Pla1-Red1 |
| --- | --- | --- | --- |
| **Data collection** |  |  |  |
| Beamline | PETRA III, P13 DESY | PETRA III, P13 DESY | PETRA III, P14 DESY |
| Wavelength (Å) | 0.9762 | 0.9762 | 0.9801 |
| Resolution range (Å) | 77.68 - 1.9<br>(1.968 - 1.9) | 57.35 - 2.599<br>(2.692 - 2.599) | 98.02 - 2.805<br>(2.905 - 2.805) |
| Space group | C121 | P1 | P2<sub>1</sub>2<sub>1</sub>2 |
| Unit cell dimensions<br>a, b, c (Å)<br>α, β, γ (°) | 122.06 66.75 88.12<br>90 118.175 90 | 71.79 72.48 73.21<br>99.963 105.702<br>118.833 | 128.644 151.337 65.408<br>90 90 90 |
| Total reflections | 333368 (32959) | 119702 (11522) | 141478 (12446)<br>90056 (3985)* |
| Unique reflections | 49226 (4854) | 34690 (3409) | 31789 (654)<br>20986 (1049)* |
| Multiplicity | 6.8 (6.8) | 3.5 (3.4) | 4.3 (3.8)* |
| Completeness (%) | 99.73 (99.02) | 97.04 (95.57) | 63.95 (20.67)<br>92.4 (96.7)* |
| Mean I/sigma(I) | 11.87 (1.53) | 7.26 (0.89) | 9.91 (0.64)<br>12.8 (2.6)* |
| R<sub>merge</sub> | 0.1145 (1.18) | 0.1545 (1.402) | 0.065 (0.479)* |
| R<sub>meas</sub> | 0.1242 (1.279) | 0.1837 (1.67) | 0.074 (0.555)* |
| R<sub>pim</sub> | 0.04764 (0.4875) | 0.09824 (0.8945) | 0.036 (0.275)* |
| CC<sub>1/2</sub> | 0.998 (0.663) | 0.99 (0.298) | 0.999(0.868)* |
| **Refinement** |  |  |  |
| Reflections used in refinement | 49223 (4854) | 34678 (3409) | 20527 (654) |
| Reflections used for R<sub>free</sub> | 2462 (243) | 1733 (171) | 1999 (63) |
| R<sub>work</sub> | 0.1690 (0.3095) | 0.2161 (0.3710) | 0.2413 (0.3401) |
| R<sub>free</sub> | 0.1923 (0.3453) | 0.2575 (0.4146) | 0.2840 (0.4666) |
| Number of non-hydrogen atoms | 4718 | 8199 | 8531 |
| Macromolecules | 4185 | 8113 | 8528 |
| Ligands | 6 | - | - |
| Solvent | 527 | 86 | 3 |
| R.M.S deviations |  |  |  |
| Bond lengths (Å) | 0.003 | 0.002 | 0.002 |
| Bond angles (°) | 0.56 | 0.51 | 0.49 |
| Ramachandran plot |  |  |  |
| Most favored (%) | 97.44 | 97.60 | 93.71 |
| Allowed (%) | 2.37 | 2.30 | 5.71 |
| Outliers (%) | 0.20 | 0.10 | 0.57 |
| Rotamer outliers (%) | 0.65 | 1.25 | 1.71 |
| Clashscore | 6.52 | 3.80 | 4.44 |
| Average B-factor | 32.65 | 64.81 | 47.93 |
| Macromolecules | 31.75 | 64.91 | 47.93 |
| Ligands | 36.74 | - | - |
| Solvent | 39.69 | 55.47 | 41.78 |

Statistics for the highest-resolution shell are shown in parentheses.
\*Statistics reported for ellipsoidal diffraction cut-off.

Our crystal structure of the Pla1-Red1 complex shows that Red1 contacts the C-terminal RRM domain of Pla1 (**Figure 2B**). The interaction between Pla1 and Red1 buries a total solvent accessible area of 1338.5 Å^2^. The interaction interface of Red1 can be largely divided into two regions. The first half comprises a short α-helix at the N-terminus (α0) that helps orient Red1 onto Pla1 (**Figure 2C**). A network of aromatic stacking interactions occurs between Pla1_RRM_ Phe468 and Red1 Trp298 and Phe305 (**Figure 2C, Supplementary Figure S5C**). This cluster is further stabilized by hydrophobic stacking of the aliphatic sidechain of Lys368, a hydrogen bond between its side chain and backbone carbonyl of Trp298, a salt bridge formed between Glu416 and Lys294 and packing of Asp513 against Phe305. Importantly, Lys368 positioned close to the C-terminus of Red1 helix α0 reads out the negative dipole of α0 and therefore further stabilizes it. The second half of Red1 is largely involved in mainchain-sidechain interactions between Red1 residues Gly306, Ser308, Asn311 and Val312 that pack against Pla1 residues Gln420, Val518, Ile520 and Asn522 (**Figure 2D**). In addition, hydrogen bonding interactions occur between Red1 Ser307 and Ser308 side chains with Pla1 Ser424 and Lys506. Interestingly, Red1 Val312-Ile314 form a β-strand (β0) that packs against Pla1 Arg491-Asp493 that also form a short β-strand, leading to the formation of an antiparallel β-zipper^50^ (**Figure 2D**). Importantly, this short β-strand is not formed in our Pla1 apo structures. Therefore, this β-strand addition by means of a β-zipper formation stabilizes and holds the second half of Red1 in position.

To provide support for the structure of Pla1-Red1 complex and validate its conformation in solution, we recorded SAXS experiments (**Supplementary Figure S5D**). The Kratky plot of the Pla1-Red1 complex shows a bell-shaped curve characteristic of well-folded globular molecules (**Supplementary Figure S5E**). The pairwise distribution curve also shows that the Pla1-Red1 complex has a globular architecture with a *D*_max_ of 10.7 nm (**Supplementary Figure S5F**). Since the flexible Pla1_RRM_ loops L1 and L2, and both Pla1 and Red1 C-termini are missing from the crystal structure, we used CORAL^51^ to first model these regions based on the SAXS data (**Supplementary Figure S5G**). Subsequent fitting of the experimental SAXS scattering profile of the complex with the crystal structure show that the data are in good agreement with a *χ*^2^ fitting-value of 1.23 (**Supplementary Figure S5H**).

### Structure-based mutation analysis *in vitro*

Based on the NMR data and our crystal structure of the Pla1-Red1 complex, we performed mutational analyses and evaluated the importance of the affected contacts in ITC measurements. We first mutated the aromatic residues in the Red1 N-terminal helix α0 by replacing them with alanine residues to create the double mutant W298/F305A. This double mutant led to a complete disruption of Pla1-Red1 interaction (**Figure 3A**). This is not surprising as stacking interactions of Red1 helix α0 residues Trp298 and Phe305 with Pla1_RRM_ Phe468 and Lys368 creates a hydrophobic core which helps orient the Red1 peptide onto Pla1 (**Figure 2C**). A charge reversal mutation of Lys368 (K368E) which reads the negative dipole moment of Red1 helix α0 also leads to a ∼13-fold loss in affinity (**Figure 3B**). Furthermore, mutation of Red1 Val313, which is part of β0-strand, to an arginine residue also leads to ∼13-fold loss in affinity (**Figure 3C**), signifying the affinity contribution of the β-zipper. In contrast, alanine replacements of Ser308 and Asn311 did not affect the interaction (**Supplementary Figure S6A, B**).

**Figure 3.**
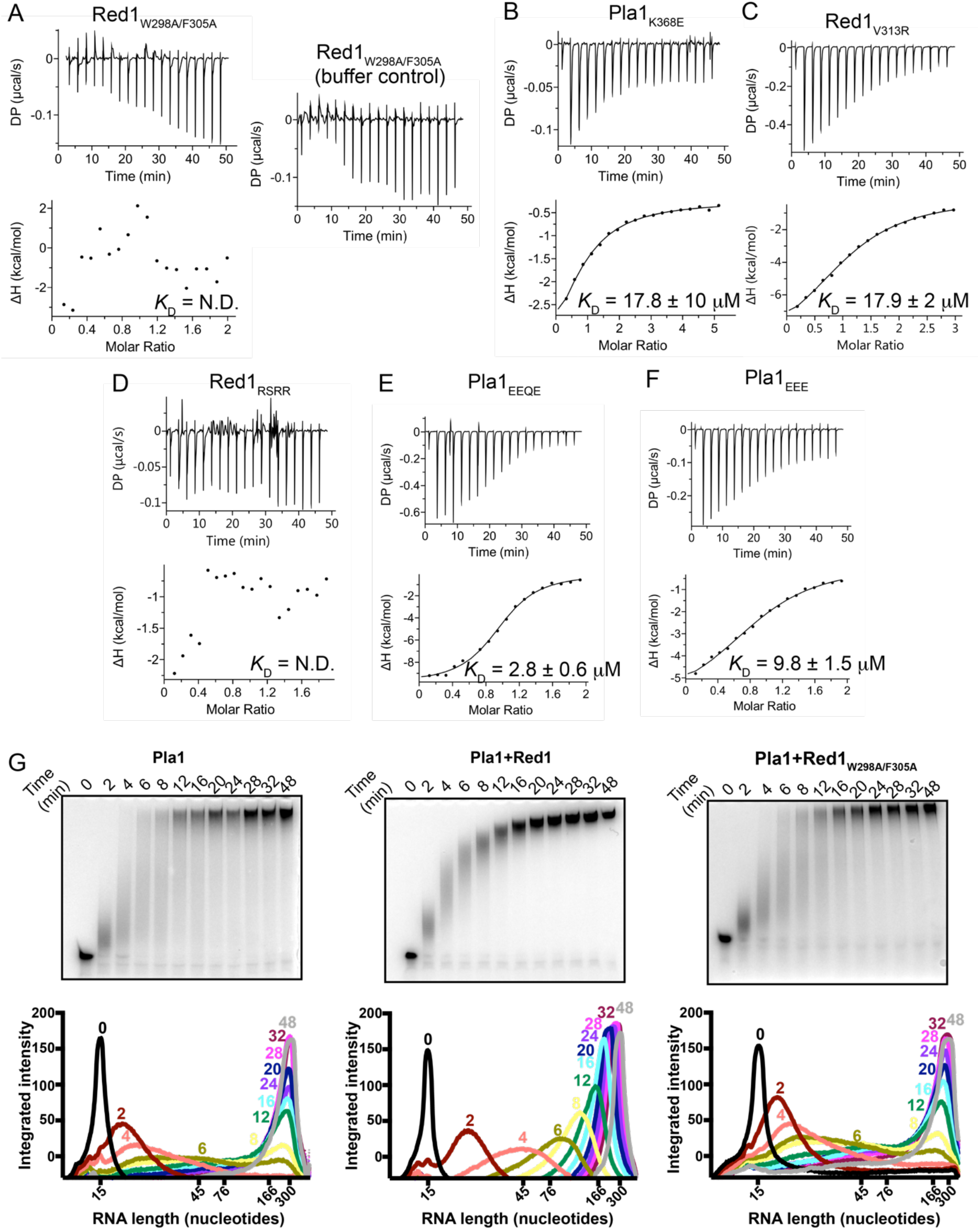
Mutational analyses of Pla1-Red1 interaction Isothermal titration calorimetry experiments with specific Pla1 and Red1 point mutants that affect their interaction interface are shown in panels. (A)-(F). The calculated dissociation constant (*K*_D_) from an average of two independent measurements is shown. (G) *In vitro* polyadenylation assay. Polyadenylation of a 5’-Cy3 labeled A15 RNA primer by Pla1 alone (left panel), in the presence of Red1_288-345_ (middle panel) and Red1_W298A/F305A_ (right panel), analyzed by 14% denaturing urea PAGE at different time points. Densitometric analyses of the gels are plotted below, where the RNA length is marked based on an RNA ladder.

Our NMR titrations had indicated that the negatively charged patch at the Red12_88-322_ C-terminus (namely residues Asp317, Ser318, Asp319 and Asp320) also show line broadening upon addition of Pla1_RRM_ (**Figure 1D, Supplementary Figure S2B**). In our crystal structure, however, these residues do not show contacts with Pla1. To probe the importance of these residues for Pla1-Red1 interaction, we made charge reversal mutations of the aspartate residues to arginine (D317/S318/D319/D320 to RSRR, named Red1_RSRR_). To our surprise this triple mutant also completely abrogated Pla1-Red1 binding (**Figure 3D**). Since 25 residues at the Pla1 C-terminus are disordered in our crystal structure and the Pla1 and Red1 C-termini are in close proximity to each other (**Supplementary Figure S6C**), we reasoned that two positively charged patches present in the disordered Pla1 C-terminus could possibly be involved in this interaction. Therefore, we created two Pla1_RRM_ charge reversal mutants: K544/K545/R546 to EEE and K559/R560/Q561/K562 to EEQE, named Pla1_EEE_ and Pla1_EEQE_, respectively. While the Pla1_EEQE_ mutant led to a ∼3-fold loss in affinity (**Figure 3E**), the Pla1_EEE_ mutant led to an ∼11-fold loss in affinity (**Figure 3F**), indicating that these charged patches of Pla1 and Red1 C-termini are in close proximity in solution and are involved in electrostatic interactions.

To assess if recombinantly purified Pla1 is active *in vitro* and if its polyadenylation activity is affected by Red1, we used *in vitro* polyadenylation assays. Recombinantly purified Pla1 was incubated with 5’-Cy3 labeled A15 RNA primer in the absence or presence of Red1 and the reaction was started by addition of ATP. As previously reported, Pla1 is active in *in vitro* polyadenylation assays^52^ (**Figure 3G**) compared to a control where the catalytic mutant of Pla1 (D153A) did not show any polyadenylation activity (**Supplementary Figure S6D**). Remarkably, Red1 affects the processivity of Pla1 and the poly(A)-tail synthesis becomes more distributive, as evidenced by the gradual increase in length of poly(A) of all products compared to a significant difference in the tail lengths of the different RNA species in the absence of Red1. This effect of Red1 on the Pla1 catalytic activity is similar to that observed for the *S. cerevisiae* homologs Pap1 and Fip1^44, 53^. Of note, the distribution of poly(A)-tail lengths of products was similar to wild-type Pla1 when the Pla1-Red1 interaction mutant (Red1_W298A/F305A_) was used (**Figure 3G**), indicating that Red1_W298A/F305A_ is unable to bind and therefore influence the processivity of Pla1.

### *In vivo* analysis of a Pla1 ‘truncated’ MTREC complex

To assess the effects of Pla1-Red1 interaction mutants, we first performed Y2H experiments using the full-length Red1 containing the W298A/F305A double mutation, together with full-length Pla1 (**Figure 4A**). Indeed, these point mutations strongly impaired the Red1-Pla1 interaction, as evidenced by the lack of colonies on -Leu -Trp -Ade media, although yeast colonies on the more sensitive -Leu -Trp -His media indicated a residual binding. However, this interaction was completely abrogated when the Red1 residues 288-345 were deleted (Red1Δ288-345) (**Figure 4A**).

**Figure 4.**
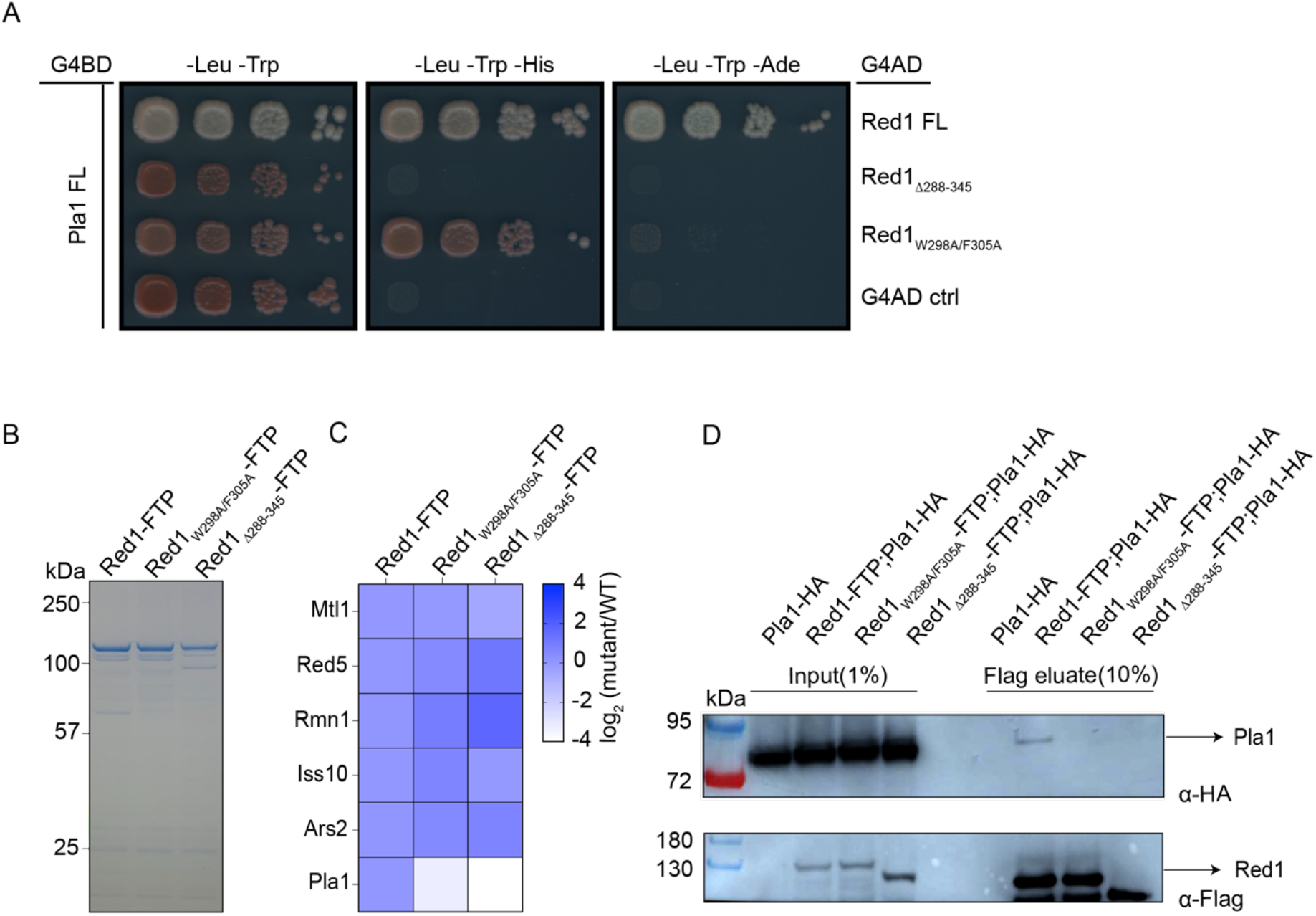
*In vivo* analysis of Pla1 ‘truncated’ MTREC complex. (A) Y2H experiments showing the effects of deletion of Red1 residues 288-345 or Red1_W298A/F305A_ double mutant on Pla1-Red1 interaction in the context of full-length proteins. (B) Coomassie blue stained SDS polyacrylamide gel of tandem affinity-purified MTREC complex using Red1 as bait, from respective mutant strains; and (C) heat map representation of mass-spectrometry (MS) analysis of these purifications. Heat maps show the changes in the amount of co-purifying MTREC proteins interacting directly with Red1 in indicated mutant strains compared to WT. Co-purifying protein amounts were normalized to the corresponding purified bait protein (WT or mutant Red1) amount. (D) Co-Immunoprecipitation (co-IP) of HA-tagged Pla1 (Pla1-HA) using Red1 as bait in WT and Red1 mutant strains. Inputs and final eluates of the tandem affinity purifications of the indicated strains were probed with α-HA to detect co-purifying Pla1-HA (top panel) and with α-Flag to detect the bait protein Red1-FTP (bottom panel).

To confirm that the MTREC complex lost its association with its Pla1 subunit in the *Red1_W298A/F305A_* and *red1_Δ288-345_* strains *in vivo*, we replaced the genomic copy of Red1 in *S. pombe* cells with a C-terminally Flag-ProtA-tagged version of wild type (WT) Red1, *Red1_W298A/F305A_* double mutant and *red1_Δ288-345_* deletion mutant. We performed tandem affinity purifications with these strains (**Figure 4B**) and analysed the final eluates using mass spectroscopy with TMT 10plex mass tag labeling (**Figure 4C**). These analyses confirmed that both Red1_W298A/F305A_ and Red1_Δ288-345_ proteins maintained their association with MTREC components, purifying similar amounts of MTREC subunits to WT Red1, with the exception of the Pla1 subunit, which was not detectable in the purification of Red1 mutants (**Figure 4C**). To further confirm that the Pla1-Red1 interaction was disrupted in the *Red1_W298A/F305A_* and *red1_Δ288-345_* mutants, we tagged the genomic copy of Pla1 with an HA-tag in WT and mutant Red1-FTP strain backgrounds and performed co-immunoprecipitation (co-IP) experiments. Western blot (WB) results showed that Pla1-HA signal could not be detected in the final eluate of our Red1-Pla1 interaction mutant strains (**Figure 4D**). Taken together, these data confirm that *Red1_W298A/F305A_* and *red1_Δ288-345_* mutants lead to a ‘truncated MTREC’, devoid of Pla1, both *in vitro* and *in vivo*.

### Pla1 activity as part of MTREC complex is required for the efficient degradation of PROMPTs

To understand the role of Pla1 in the recognition and degradation of CUTs, we sequenced poly(A)^+^ RNA from WT, Red1 knock-out (*red1*Δ), *Red1_W298A/F305A_* and *red1_Δ288-345_* strains. Metagene plots of sense and antisense RNA levels 500 bp upstream and downstream of all *S. pombe* genes (**Supplementary Figure S7A,B**) and a subset of genes (2400 genes, ∼40% of all genes) filtered for detectable levels of PROMPTs (**Figure 5A-C**) in WT and red1 mutant cells confirmed the strong accumulation of PROMPTs, antisense (AS) RNAs and 3’ intergenic transcripts (3’IGTs) in *red1Δ* cells compared to WT. Interestingly, *Red1_W298A/F305A_* and *red1_Δ288-345_* cells showed only a moderate but highly reproducible accumulation in the levels of PROMPTs, while AS RNAs and 3’ intergenic transcript (3’IGT) levels were not affected. Meiotic genes and levels of intronic sequences were also unaffected in these mutants, compared to WT (**Supplementary Figure S7C-F**). To exclude the possibility that changes in the Red1-Pla1 interaction mutants remained undetected due to the complete lack of poly(A) tails in stabilized transcripts, we repeated these analyses, using total RNA sequencing (**Supplementary Figure S8A, B)**. While the total RNA sequencing detected less antisense RNA transcripts in general, the difference between WT and Red1-Pla1 interaction mutants was nearly identical to the poly(A)^+^ transcriptome analysis, strongly suggesting that the role of Pla1 in the context of the MTREC complex is not the initial poly-adenylation but rather the extension of the poly(A) tails of MTREC substrates, leading to the well documented hyper-adenylation of these transcripts^33–35^.

**Figure 5.**
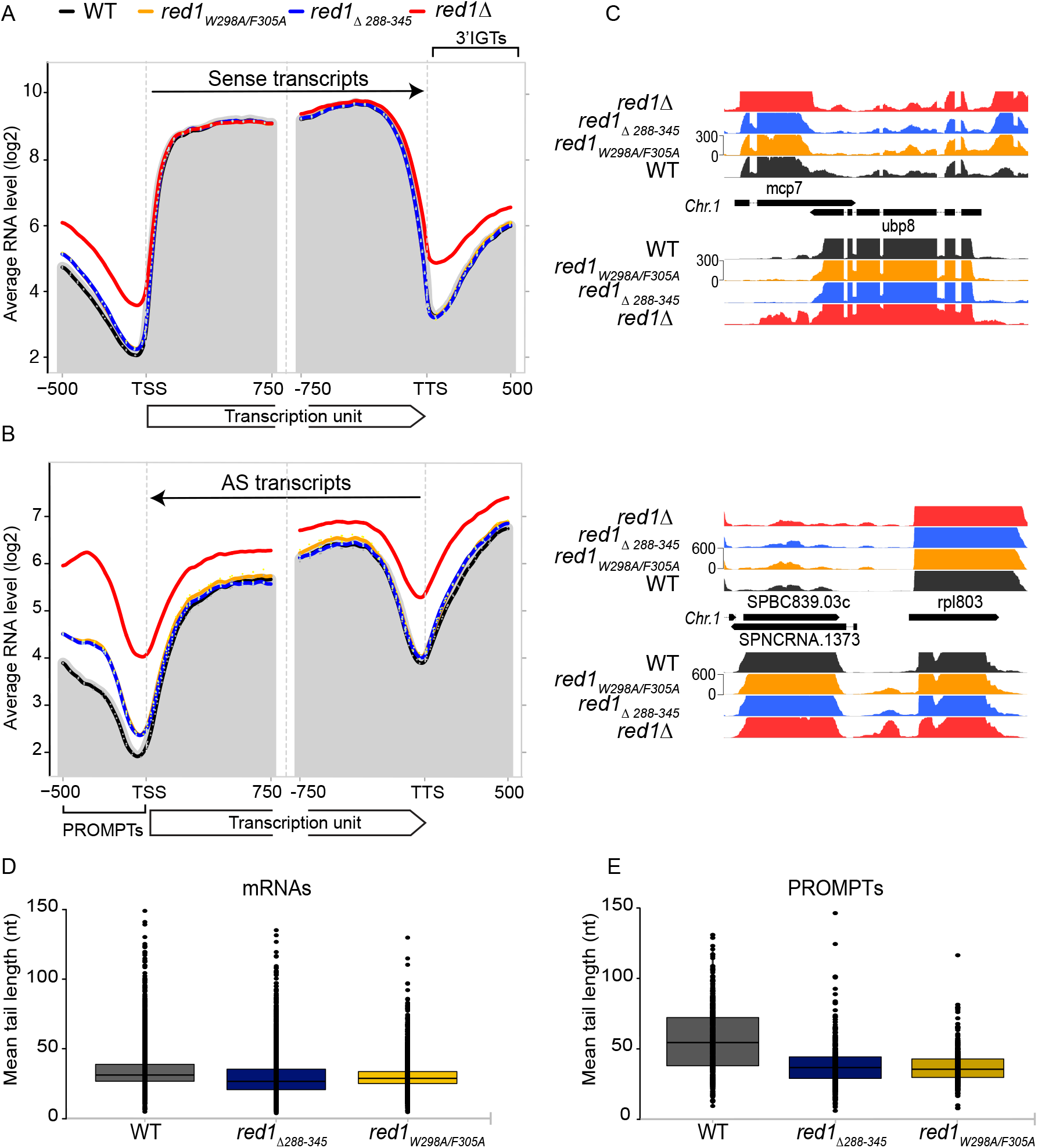
Pla1 in the context of the MTREC complex is responsible for degradation of PROMPTs. (A and B) Metagene profile of sense (A) and antisense (B) RNA levels in the indicated strains for a subset of *S. pombe* genes with detectable levels of PROMPTs (2400 genes). The geometric average of RNA levels from 500bp upstream to 750bp downstream of the transcription start site (TSS) and 750bp upstream to 500bp downstream of the transcription termination site (TTS) are shown. Solid lines represent the average of two replicates for all indicated strains, with the exception of *red1Δ* which represent a single dataset. Dotted lines indicate the individual biological replicates. The grey shading represents the average RNA levels in the WT strain. (C) Strand-specific RNA-seq read coverage of a representative set of genes in WT and mutant strains. (D and E) Box-plot of mean poly(A) tail length distribution of mRNAs (D) and PROMPTs (E) for the WT and indicated mutant strains. Dots represent the mean poly(A) tail length of individual mRNAs/PROMPTs, boxes show the 25-75 percentile range and the lines represent the median values.

### Disrupting Red1-Pla1 interaction affects the poly(A) tail lengths of CUTs and meiotic mRNAs

To assess poly(A)-tail length of CUTs and meiotic mRNAs in WT and Red1-Pla1 interaction mutants, we used Oxford Nanopore Technologies’ direct RNA sequencing technology, combined with the *tailfindr* R package^54^. This package can estimate poly(A)-tail length of individual reads from the length of the monotonous low-variance raw signal, corresponding to poly(A)-tails, at the beginning of each read and combines this information with the unique read ID. After mapping the individual reads to the *S. pombe* genome, we can assign the poly(A) tail length information to these mapped reads, determine mean poly(A)-tail length of selected transcripts (e.g., all sequenced transcripts of a particular gene or a particular CUT), and display this information for selected population of transcripts (e.g., CUTs or meiotic mRNAs). Since CUTs and meiotic mRNAs are extremely low abundant in WT cells, and might represent a specific sub-population of these transcripts that escaped MTREC and exosome-mediated degradation, we decided to measure the poly(A)-tail length of RNA transcripts associated with the MTREC complex. We have previously shown that RNA immunoprecipitation (RIP) of MTREC complex components strongly enrich CUTs and meiotic mRNAs^26^, therefore, we purified Flag-ProtA-tagged WT Red1 as well as Red1_W298A/F305A_ and Red11__Δ288-345__ mutants, using conditions that preserve RNA-protein complexes and we analysed the co-precipitating RNA transcripts by direct RNA sequencing.

Previous studies evaluated the poly(A)-tail length of individual transcripts (individual meiotic mRNAs or individual CUTs^26, 33–35^) and concluded that these transcripts are hyper-adenylated, compared to normal mRNAs. Our genome-wide data confirm these findings, estimating median poly(A)-tail length of mRNAs to be around 32 nucleotides (nts) in WT *S. pombe* cells, while PROMPTs and meiotic mRNAs have nearly twice longer poly(A)-tails.

The mean poly(A)-tail length distribution of IP-ed mRNA transcripts (representing mainly contaminating RNA transcripts in these RIP experiments) shows only a minor difference between WT and mutants (**Figure 5D**; median values are 32, 29 and 27 nts in WT, *Red1_W298A/F305A_* and *Red11__Δ288-345__* mutants, respectively), while PROMPTs show poly(A)-tail length decreased by ∼20 nts in *Red1_W298A/F305A_* and *Red11__Δ288-345__* mutants, compared to WT (**Fig 5E**; median 54 in WT to median 35 and 37 nts in *Red1_W298A/F305A_* and *Red11__Δ288-345__* mutants, respectively). Interestingly, while Red1-Pla1 interaction mutants did not affect the degradation of AS RNAs or meiotic mRNAs, the median poly(A)-tail lengths of these transcripts were also reduced in the mutant strains (**Supplementary Figure S8C,D)**, although less pronounced than for PROMPTs (meiotic mRNAs: median 56 nts in WT to 32 nts in *Red1_W298A/F305A_* and 42 nts in *Red11__Δ288-345__* mutants; AS RNAs: median 41 nts in WT to 32 nts in *Red1_W298A/F305A_* and 35 nts in *Red11__Δ288-345__* mutants). Overall, these experiments show that the poly(A)-tail length of CUTs and meiotic mRNAs are reduced by ∼20 nts in *Red1_W298A/F305A_* and *Red11__Δ288-345__* mutants, while the poly(A)-tail length of mRNAs remains unaffected.

### MTREC-Pla1 interaction is required for heterochromatic island formation at meiotic genes

Recent studies reported the involvement of the CPF, including Pla1, in the formation of small, facultative heterochromatic islands at meiotic genes^55, 56^. We wondered if the Red1-Pla1 interaction might play a direct role in this process. We carried out ChIPseq experiments to detect H3K9me2 modifications throughout the fission yeast genome in WT, *red1Δ* and the Red1-Pla1 interaction mutant strains (*Red1_W298A/F305A_ and Red11__Δ288-345__*). We used 3 independent WT strains in this experiment, with slightly different genetic backgrounds (P1, P419, F3230; see **Supplementary Table S3** for genotype information), to account for potential variabilities in the appearance of these facultative heterochromatic islands. We detected all previously described heterochromatic islands within the *S. pombe* genome^57–60^, and their size and H3K9me2 enrichment levels were remarkably uniform between the 3 independent WT strains (**Figure 6A, B**). We also confirmed that most of the facultative heterochromatic islands, mainly located at meiotic genes, are dependent on an intact MTREC complex, as evidenced by the complete absence of these islands in the *red1Δ* cells (**Figure 6B, C** and ^55, 56^). Other islands (non-meiotic islands) are independent of the MTREC complex and they are unaffected in the *red1Δ* strain^57^ (**Figure 6D**). Interestingly, H3K9me2 enrichment levels at all MTREC-dependent facultative heterochromatic islands were strongly reduced in the Red1-Pla1 interaction mutants, while MTREC-independent islands and the major heterochromatic regions (pericentromeric regions, telomers, mating-type and rDNA-region) were unaffected (**Figure 6B-D**). The reduction of the H3K9me2 levels at these islands were comparable to the effect that was reported in CPF mutant strains^56^. This finding is remarkable, since *Red1_W298A/F305A_* and *Red11__Δ288-345__* mutants do not interfere with MTREC-mediated degradation of meiotic mRNAs. These results suggest that while the recruitment of MTREC complex to meiotic mRNAs does not require Pla1 interaction with the MTREC complex, efficient establishment and/or maintenance of the heterochromatic islands at these loci is dependent on the intact MTREC-Pla1 physical interaction.

**Figure 6.**
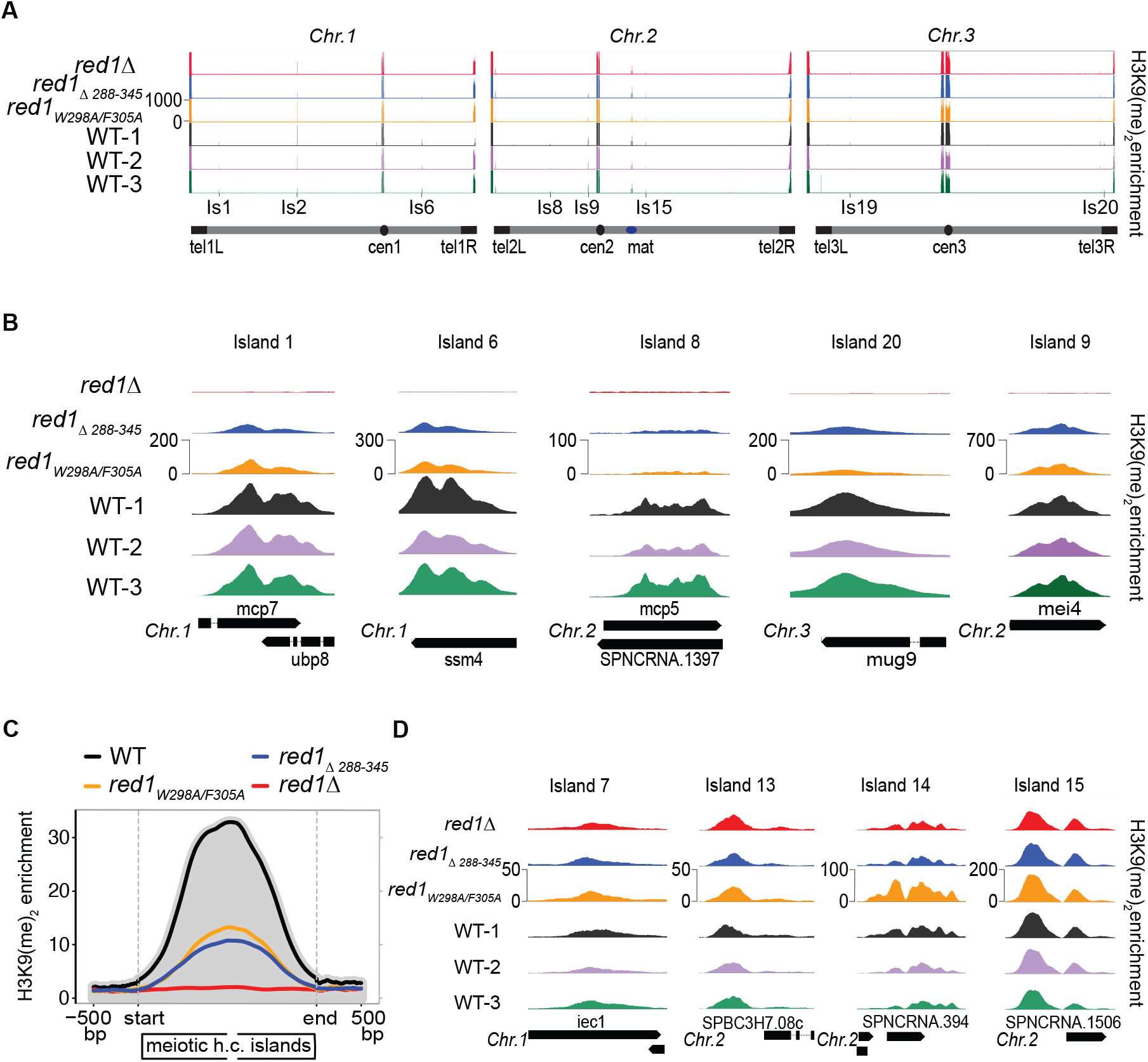
Pla1 – MTREC interaction is required for efficient assembly of facultative heterochromatic islands at meiotic genes. **(A)** Genome-wide H3K9me2 ChIP-seq analysis of 3 independent WT strains (WT-1 is the isogenic WT strain for the mutants) and red11β, Red1_W298A/F305A_ and Red11__Δ288-345__ mutants, showing the H3K9me2 enrichment levels (scale 0-100) for the 3 chromosomes of the *S. pombe* genome (length not to scale). Centromeres, Mating-type locus, telomeres and a selection of facultative heterochromatic islands are indicated. **(B)** Representative examples of *red1Δ*-sensitive meiotic heterochromatic islands in WTs and mutant strains are shown in higher resolution (scales are indicated for individual islands). **(C)** Average of H3K9me2 enrichment over all meiotic heterochromatic islands (12 islands: is1(mcp7); is1.5(tht2); is1.6(SPAC631.02), is2(mug8); is4(SPAC8C9.04/SPNCRNA.925); is5(vps29); is6(ssm4); is8(mcp5); is9(mei4); is16(mbx2/SPNCRNA.1626); is17(mug45); is20(mug9)) plotted on a linear scale. The plots represent the geometric average of enrichment values from 500bp upstream to 500 bp downstream of the islands, with the island regions scaled to the same lengths for all islands (stretched or condensed to 2000bp). The grey shading represents the average H3K9me2 enrichment levels in the WT strain. **(D)** Representative examples of non-meiotic (*red1Δ*-insensitive) heterochromatic islands in WTs and mutant strains (scales are indicated for individual islands).

### Interaction of MTREC complex with CPF

3’-end processing of pre-mRNA involves the complex assembly and action of a number of proteins, including the highly conserved CPF^61–64^. CPF is involved in site-specific endonucleolytic cleavage of the pre-mRNA followed by addition of a poly(A)-tail at its 3’ end, which is required for nuclear export of mRNAs. In *S. cerevisiae*, the CPF is composed of three enzymatically active modules of which the polymerase module encompasses, among others, the Pap1^64^ and Fip1 subunits, and Fip1 has been suggested to tether Pap1 to CPF^44, 45, 65–67^. Consistently, the interaction between Fip1 and PAP is also conserved in humans^47^. Fip1 is an intrinsically disordered protein with short regions directly contacting Pap1/Yth1^65–67^. The crystal structure of Pap1 in complex with a 26 amino acid Fip1 peptide shows that the N-terminus of the short peptide forms a parallel β-ribbon with Pap1 and adds an antiparallel β-strand extending the β-sheet interface of Pap1^65^. Structure alignment of the Pap1-Fip1 complex with our Pla1-Red1 structure shows that Fip1 and Red1 partially occupy a similar binding interface on the polymerase (**Supplementary Figure S9A**). Interestingly, there have been suggestions that the MTREC complex and 3’-end processing machinery act in a coordinated fashion where the CPF complex cleaves the RNA destined for degradation, thereby creating an entry point for the MTREC complex^55, 56^.

Given the existence of an overlapping binding interface of Fip1 and Red1 on the polymerase, we wondered if a direct competition between Fip1 and Red1 for binding to Pla1 exists. Based on sequence alignment between Fip1 and its fission yeast ortholog Iss1 and the extended Pap1 binding site recently identified within Fip1^67^ (**Supplementary Figure S9B**), we tested the binding of a 46-amino acid construct of Iss1 (residues 30-76) to Pla1 using ITC. We found that Iss1 binds to Pla1 with micromolar affinity (*K*_D_ = 2.5 ± 0.16 µM) (**Figure 7A**). In competition experiments, where Red1_288-345_ is titrated into a pre-formed Pla1_RRM_-Iss1 complex, we observed an ∼8-fold decrease in affinity of Pla1 for Red1 (*K*_D_ = 11 ± 1.6 µM) (**Figure 7B**). In contrast, a titration of Iss1 into Pla1_RRM_-Red1_288-345_ complex is unable to displace Red1 from the complex (**Figure 7C**). These data show the existence of negative cooperativity between Iss1 and Red1 due to an overlapping binding interface on Pla1_RRM_. The inability of Iss1 to outcompete Red1 from the Pla1-Red1 complex, even though it binds only ∼1.8-fold weaker compared to Red1_288-345_, prompted us to investigate the binding kinetics of the two proteins. We therefore used bio-layer interferometry (BLI) to measure possible differences in the *K*_on_/*K*_off_ rates of biotinylated Red1_288-345_ (**Figure 7D**) or Iss1 (**Figure 7E**) immobilized on streptavidin-coated biosensors to Pla1_FL_. Compared to Iss1 (*k*_on_ = 2.1*10^5^ M^-1^s^-1^, *k*_off_ = 2.4*10^-2^ s^-1^), Red1_288-345_ has a faster *k*_on_ (2.8*10^5^ M^-1^s^-1^) and a slower *k*_off_ (1.6*10^-2^ s^-1^) (**Supplementary Table S2**).

Furthermore, we have previously observed that a member of accessory cleavage factors (CF) of CPF known as Msi2 (Hrp1 or CF IB in *S. cerevisiae*), co-purifies with MTREC^26^, suggesting a possible association between these two machineries. We therefore wanted to understand and identify whether Msi2 could directly interact with individual components of MTREC. Indeed, we observed that Msi2 binds to two distinct members of the MTREC complex, Pab2 and Mmi1, in Y2H experiments (**Figure 7F, Supplementary Figure S9C**). In summary, these data show that multiple points of association exist between CPF and the MTREC complex and suggest a close functional cooperation between these two complexes in RNA surveillance.

**Figure 7.**
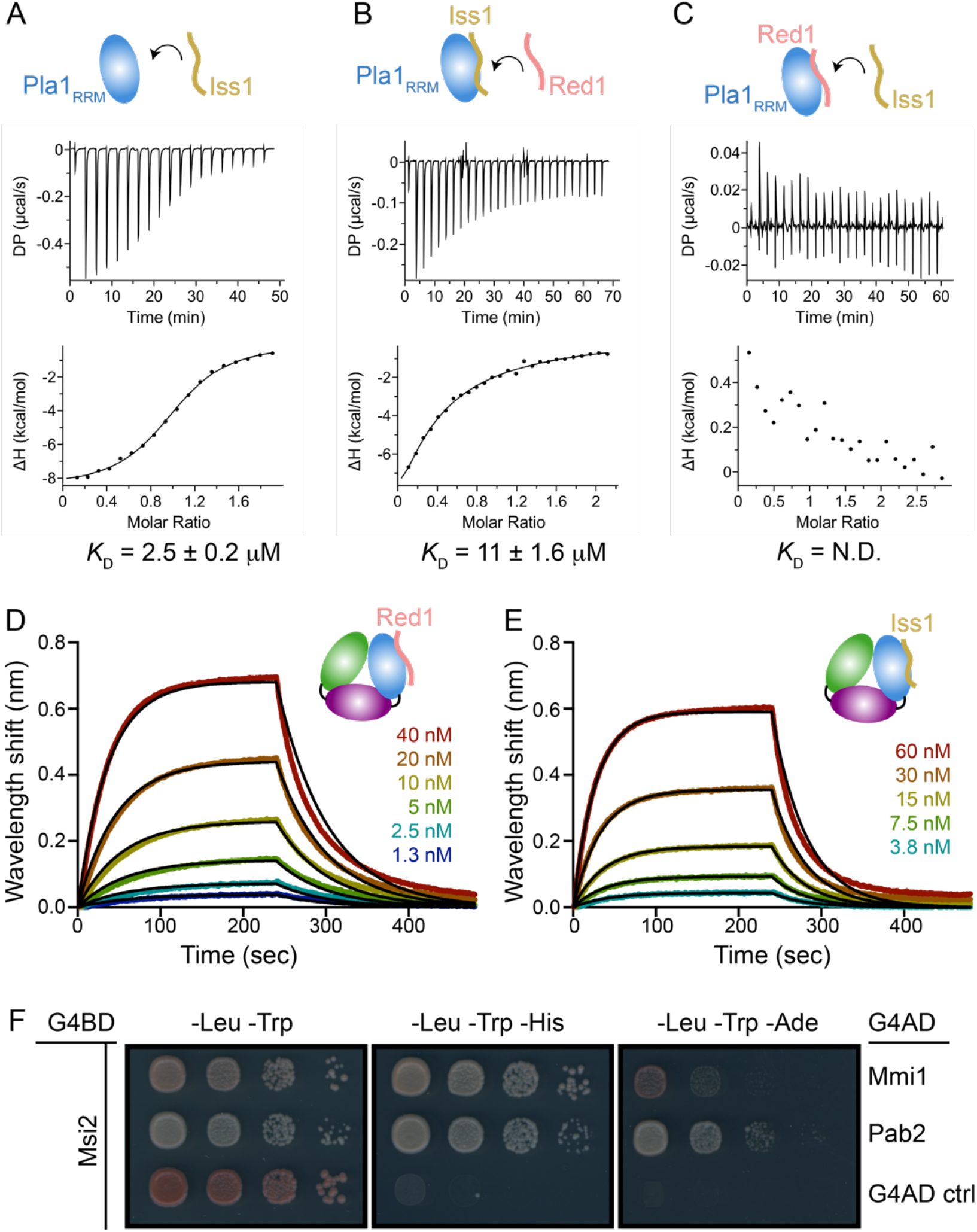
Interactions between MTREC and CPF. (A) ITC titration of Iss1 into Pla1_RRM_ domain. (B) ITC titration of Red1_288-345_ into a pre-formed complex of Pla1_RRM_-Iss1. (C) ITC titration of Iss1 into a pre-formed complex of Pla1_RRM_-Red1_288-345_. The calculated dissociation constants (*K*_D_) from an average of two independent measurements are shown. BLI kinetic analyses of biotinylated Red1_288-345_ and Iss1 with varying concentrations of Pla1_FL_ are shown in panels (D) and (E), respectively. Black lines represent fits to the experimental data obtained using a 1:1 binding global fitting model. (F) Y2H experiments show interaction between Msi2 fused to G4BD and MTREC components Mmi1 or Pab2 fused to G4AD compared to an auto-activation control (ctrl).

## Discussion

The multi-subunit MTREC complex serves as the exosome-adaptor complex required for degradation of CUTs in *S. pombe*. In this study, we identified and characterised the Pla1-Red1 interaction using Y2H, NMR and X-ray crystallography. We showed that the C-terminal RRM domain of Pla1 binds to a 58-residue region of the MTREC core component Red1 (residues 288-345), tethering it to the MTREC complex (**Figure 1**). Our crystal structure of the Pla1-Red1 complex showed that Red1 is largely unstructured but comprises an α-helix and a β-strand at the N- and C-termini, which hold Red1 in position to form a tight binding interface with Pla1_RRM_ (**Figure 2B**). Using structure-based mutational analyses, we identified a Red1_W298A/F305A_ double mutant in the N-terminal α-helix that severely compromises the Pla1-Red1 interaction *in vitro* (**Figure 3A**) and *in vivo* (**Figure 4**), while deletion of the entire interaction surface in the Red11__Δ288-345__ mutant leads to no detectable interaction between the MTREC complex and Pla1. Using these Pla1-Red1 interaction mutants, we showed that the truncated MTREC, devoid of Pla1, is unable to hyper-adenylate CUTs and meiotic mRNAs, leading to the inefficient degradation of these transcripts. In addition, the Pla1-Red1 interaction is required for the efficient establishment and/or maintenance of facultative heterochromatic islands around meiotic genes, as evidenced by the strongly impaired levels of H3K9me2 at these loci in the mutants.

Interestingly, similar to the MTREC complex where Pla1 is tethered to the complex via Red1, the *S. cerevisiae* homologue of Pla1 (Pap1) has been shown to be flexibly tethered to the CPF core machinery via the intrinsically disordered protein Fip1^65–67^. Structural superimposition of Pla1-Red1 and the *S. cerevisiae* Pap1-Fip1 complexes show that Fip1 and Red1 occupy a similar binding interface on the respective polymerase (**Supplementary Figure S9A**). Surprisingly, Fip1 binds to Pap1 with picomolar affinity^65^, while in our experiments the *S. pombe* homologue of Fip1 (Iss1) binds with a weaker, low micromolar affinity (*K*_D_ = 2.5 ± 0.16 µM, **Figure 7A**), which can be attributed to the low sequence conservation of Fip1/Iss1 in the polymerase binding region and different assay conditions (^67^, **Supplementary Figure S9B**). In our *in vitro* competition experiments, we observed that Red1 is able to outcompete Iss1 from its complex with Pla1 (**Figure 7B, C**) owing to faster *k*_on_ and slower *k*_off_ rates of Red1 when compared to that of Iss1 (**Figure 7D, E and Supplementary Table S2**). While these results might differ in the context of the complete CPF and MTREC complexes, there is no evidence to suggest additional interactions between Pap1/Pla1 and other components of CPF or MTREC, besides Fip1^67^ and Red1^29^, respectively. Such a negative cooperation between Iss1 and Red1 is suggestive of Pla1 sequestration from CPF via Red1 to hyper-adenylate CUTs as part of the MTREC complex, although the simultaneous existence of separate copies of Pla1 in these complexes cannot be ruled out (**Figure 8**).

**Figure 8.**
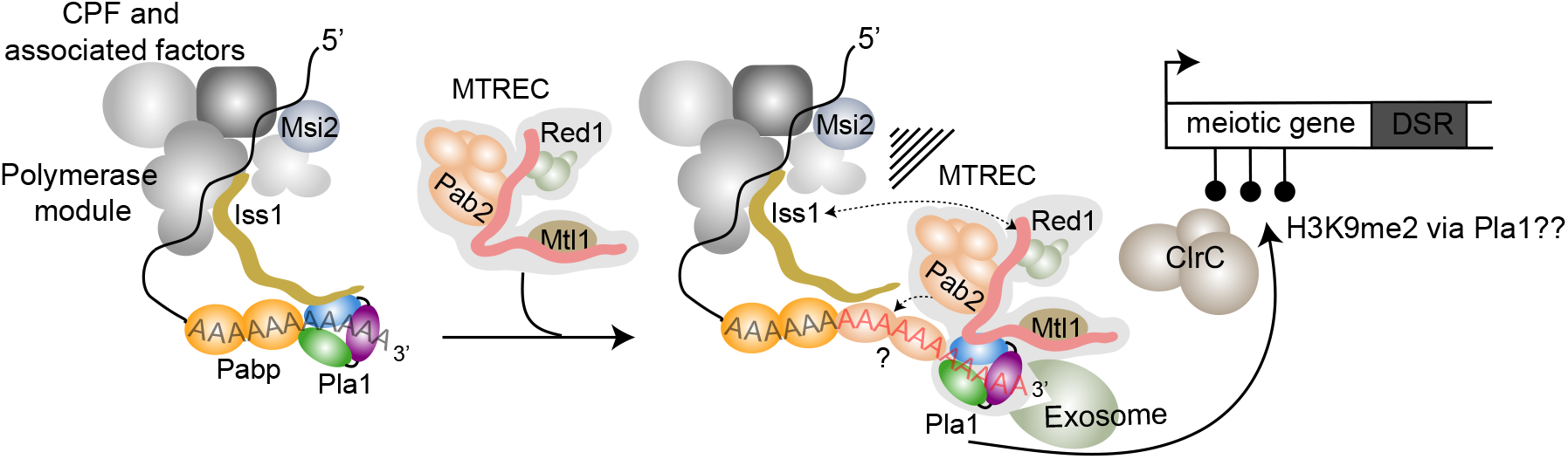
Model of the role of Pla1 in MTREC mediated degradation of CUTs. MTREC complex is recruited to CUTs and meiotic mRNAs during their transcription by a not well understood mechanism. Pla1, as part of the CPF, is responsible for the initial poly-adenylation of CUTs and the resulting poly(A)-tail is likely bound by the canonical poly(A) binding protein Pabp. MTREC complex sequesters Pla1 from CPF via Red1, replacing Iss1 that anchors Pla1 to CPF. MTREC-bound Pla1 hyper-adenylated CUTs and the non-canonical poly(A)-binding protein, Pab2 is loaded on the extended poly(A)-tail to facilitate exosome mediated degradation of CUTs. Both the CPF complex and the Red1-Pla1 interaction are required for the efficient recruitment of the ClrC complex to methylate histone H3K9 at meiotic heterochromatic islands, indicating a sophisticated functional interplay between CPF and MTREC complex during the transcriptional termination, end-processing and degradation of CUTs.

Our total RNA sequencing experiments that don’t require any poly(A)-tail to detect transcripts, did not uncover transcripts without poly(A)-tail in the Pla1-Red1 interaction mutants, strongly suggesting that the role of Pla1 in the context of the MTREC complex is not the initial poly-adenylation of CUTs, but rather the extension of the poly(A) tail of these transcripts. We carried out poly(A)-tail length analyses and showed that the median poly(A)-tail length of mRNAs is ∼32 nts in WT *S. pombe* cells, while PROMPTs and meiotic mRNAs harbour a poly(A)-tail of about 20 nts longer. The median poly(A)-tail length of PROMPTs and meiotic mRNAs were 54 and 56 nts, respectively, while AS RNAs showed a somewhat shorter median tail length of 41 nts. It is unclear if PROMPTs and meiotic mRNAs are indeed more extensively hyper-adenylated than other subclasses of CUTs, or whether this lower median value is a data analysis artefact. While PROMPTs and meiotic mRNAs can be annotated and bioinformatically captured relatively easily, intergenic and long antisense CUTs are harder to define and our analysis likely also includes stable ncRNAs which are not hyper-adenylated. Nevertheless, our genome-wide poly(A)-tail length analysis confirms that, in WT *S. pombe* cells, CUTs and meiotic mRNAs are hyper-adenylated, similar to previous reports that analysed individual CUTs or meiotic mRNAs^26, 33, 34, 68^. However, in the Pla1-Red1 interaction mutant strains, the poly(A)-tail lengths of CUTs and meiotic mRNAs are decreased, while the poly(A)-tails of mRNAs (other than meiotic mRNAs) are not affected. In these mutants, all RNA species, including CUTs and mRNAs, have a relatively uniform median poly(A)-tail length of ∼30 to 35 nts, likely representing the consistent action of Pla1 as part of CPF. These results further support our model that Pla1, as part of the MTREC complex, hyper-adenylates MTREC target RNAs, extending the existing poly(A)-tail by approximately 20 nts (**Figure 8**).

It is intriguing why the addition of 20-25 nt poly(A)-tails to PROMPTs leads to their efficient degradation, while PROMPTs harbouring shorter poly(A)-tails, due to decoupling of Pla1 from the MTREC complex, are inefficiently degraded by the nuclear exosome. Given that the core exosome channel leading to the Dis3 active site can accommodate ∼25-30 nts^69^, the additional hyper-adenylation of CUTs by MTREC must serve another purpose. One possibility is that hyper-adenylated CUTs are instantaneously bound by the MTREC component poly(A) binding protein 2 (Pab2), making the extended poly(A)-tail sterically unavailable for binding to Pabp/Pab1, the major poly(A) binding protein involved in nuclear export and poly(A)-tail length control in mRNAs (reviewed in ^70^). While Pab2 does not directly influence the polyadenylation activity of Pla1^52^, as opposed to its mammalian homologue PABPN1^71, 72^, it is required for efficient nuclear exosome-mediated degradation of CUTs, meiotic mRNAs and unspliced pre-mRNAs^25, 26, 68, 73, 74^.

Determination of RNA fate is an extremely complex process requiring timely protein-protein interactions. In fact, components of the NEXT complex co-purify with those of the 3’-end processing machinery, suggesting their co-occurrence at the 3’-ends of nascent RNAs^75^. However, we found that Msi2 was the only component of the 3’-end processing machinery co-purifying with MTREC complex when we use benzonase treatment, indicating direct protein-protein interactions^26^. Indeed, in our Y2H experiments, Msi2 interacts with two subunits of the MTREC complex, Mmi1 and Pab2 (**Figure 7F**). Importantly in *S. cerevisiae*, Hrp1 (homologue of Msi2) has been reported to participate in surveillance of CUTs^76^, Nrd1-dependent termination^77^ and cytoplasmic nonsense-mediated decay^78^, in addition to its canonical role in the 3’-end processing machinery^79^. Whether the interaction between Msi2 and Mmi1/Pab2 helps the recruitment of MTREC to CUTs, or MTREC-bound transcripts are cleaved and poly-adenylated by a specialised form of CPF, remains a topic for future investigations.

In recent years, new evidence is emerging for a co-transcriptional interaction between CPF and MTREC components^80^. Both the MTREC complex and multiple subunits of the CPF are required for the establishment and/or maintenance of facultative heterochromatic islands at meiotic genes^27, 33, 55, 56, 60, 81^. The fact that the Pla1-Red1 interaction mutants strongly destabilise these heterochromatic islands, very similar to the deletion of various CPF subunits, suggests that Pla1, as part of the MTREC complex, must functionally closely interact with CPF, as opposed to having only a strictly downstream function in this process. The exact nature of these interactions and the functional interplay between MTREC, CPF and the ClrC complex that is recruited to these loci to methylate histone H3K9, is not yet understood and will require further studies to delineate this fascinating interaction between the RNA surveillance machinery and the epigenetic regulation of the genome.

## Materials and Methods

### PCR and cloning

The DNA sequence encoding Pla1_FL_ (residues 1-566), Pla1_Δ14_ (residues 1-542), Pla1D153A (catalytic mutant, residues 1-566) were cloned and ligated into a modified pET24d vector containing non-cleavable N-terminal His6 tag. The C-terminal RRM domain of Pla1 (Pla1_RRM_, residues 352-566) was sub-cloned from Pla1_FL_ into a modified pET24d vector containing N-terminal His6 tag with the Tobacco-Etch-Virus site present before the corresponding protein. All wild-type and mutant proteins of Red1 with different boundaries were cloned into a modified pET24a vector containing N-terminal GB1 tag-TEV site before the corresponding protein sequence and a C-terminal non-cleavable His6 tag. Site directed mutagenesis with QuickChange Lightning kit was used to introduce point mutations in Pla1 and Red1 proteins, or deletion of residues 288-345 in Red1 and the mutations were confirmed using DNA sequencing.

### Protein expression and purification

The plasmids were transformed in BL21 (DE3) Rosetta2 chemically competent cells for Pla1, grown at 37 °C up to an OD of 1.4-1.6 in Luria broth or Terrific broth subsequently expressed at 20 °C for ∼16 hr after induction with 0.5 mM IPTG. For Red1, the plasmids were transformed in BL21 (DE3) chemically competent cells, grown at 37 °C up to an OD of 1 in auto-induction media ^82^ and subsequently expressed at 20 °C for ∼16 hr. For isotope-labeled proteins, the bacteria were grown in M9 minimal media supplemented with ^13^C-glucose and/or ^15^NH_4_Cl. Cells were lysed in IMAC buffer containing 20 mM Tris pH 7.5, 200 mM NaCl, 20 mM Imidazole and 2 mM ß-mercaptoethanol. The proteins were purified over 1-2 ml His-Trap FF columns (GE Healthcare) with elution buffer containing 250 mM Imidazole. In case of Red1 or Pla1 CTD, overnight tag-cleavage using Tobacco Etch Virus protease and simultaneous dialysis of the proteins into 20 mM HEPES pH 7.5, 150 mM NaCl, 2mM BME was carried out. The Red1 proteins were further purified over a second IMAC column where the cleaved proteins (still containing C-terminal His6 tag) were eluted using 250 mM Imidazole. Finally, the Red1 proteins were polished using gel filtration (Superdex 75 16/60) column (GE Healthcare) equilibrated with buffer containing 20 mM HEPES pH 7.5, 150 mM NaCl and 1 mM DTT. For ITC experiments, the DTT was replace by 2 mM BME and for NMR experiments the proteins were purified in buffer containing 20 mM sodium phosphate pH 6.5, 100 mM NaCl, 1mM DTT. For Pla1 proteins, after the IMAC column the proteins were diluted to buffer containing 50 mM NaCl and further purified using 1 ml Resource Q cation exchange column (GE Healthcare), where they were eluted with a linear gradient of 50 mM NaCl to 1 M NaCl. Gel filtration (Superdex 200 16/60) column (GE Healthcare) was used as a final polishing step, where the protein was purified in buffer containing 20 mM HEPES pH 7.5, 150 mM NaCl and 1 mM DTT. For crystallization of the apo form, the salt concentration was adjusted to 200 mM.

### X-ray crystallography

Pla1_FL_ crystallized at a concentration of 12.7 mg/ml in a drop containing 0.2 M lithium citrate, 20% PEG 3350 at 4 °C as needles within 1 week. Pla11β14 crystallized at a concentration of 6.3 mg/ml in a drop containing 0.2 M sodium formate, 20% PEG 3350 at 18 °C as thin plates within 12 days. The Pla1-Red1 complex was prepared by addition of 1.2-fold molar excess of Red1_288-345_ over Pla1_FL_ and subsequent purification over a size exclusion column in buffer containing 20 mM HEPES pH 7.5, 150 mM NaCl and 1 mM DTT, to remove excess Red1. The complex crystallized in a drop containing 0.2 M potassium formate, 20% PEG 3350 at 18 °C as thin plates within 3 weeks.

Crystals were flash frozen in mother liquor supplemented with 20% glycerol. Several datasets for the crystals were collected at P13 and P14 beamlines at PETRA III, EMBL Hamburg. Datasets from best diffracting crystals were then processed with XDS software package^83^ and the structure was solved by molecular replacement using PDB ID: 2HHP (chain A). The missing residues of Pla1 were built using Coot model building software^84^ with multiple rounds of model building and refinement with Phenix^85^. The Pla1_FL_ apo structure was then used as a molecular replacement model for solving the structure of Pla1-Red1 complex. The data were processed with XDS software package^83^. Since the crystal showed anisotropic diffraction, STARANISO server (http://staraniso.globalphasing.org)^86^ was used to apply an anisotropic correction to the data. The anisotropic correction provided an improvement in the overall interpretability of the map especially for the Pla1-Red1 interface residues. Multiple rounds of model building were done in Coot^84^ and refinement was performed with Phenix^85^.

### NMR spectroscopy

All spectra were recorded at 293K on Avance III Bruker NMR spectrometers with proton Larmor frequencies of 600 MHz, 700 MHz or 800 MHz, equipped with cryogenic (600 MHz, 800 MHz) or room temperature (700 MHz) triple resonance gradient probes. NMR experiments were performed with Red1 constructs (Red1_288-345_ or Red12_88-322_) harbouring a GB1-tag at the N-terminus and a His6-tag at the C-terminus. For backbone assignment of Red12_88-322_, data were recorded on 1 mM protein in buffer containing 25 mM sodium phosphate pH 6.5 and 100 mM NaCl, supplemented with 10 % D_2_O for lock. Protein backbone resonance assignments were obtained using 3D HNCA, HNCACB, CBCA(CO)NH and HNCO^87^. NMR titrations were done by adding 1.5-fold excess of Pla1_RRM_ into 0.3 mM ^15^N-labeled Red1 constructs (Red1_288-345_ or Red12_88-322_) and recording ^1^H,^15^N HSQC spectra. All experiments were performed at 293K. Spectra were processed in NMRPipe/Draw^88^ and analysed in CCPN Analysis^89^.

### Small angle X-ray scattering experiments

Measurements for Red1_288-345_ in complex with Pla1_FL_ were performed at 20 °C at EMBL P12 beamline, PETRA III (DESY, Hamburg, Germany) at concentrations ranging from 0.05-1.7 mg/ml in buffer containing 20 mM HEPES pH 7.5, 150 mM NaCl, 1mM DTT. Forty successive frames with 0.195 s/ frame were recorded using an X-ray wavelength of *λ*= 0.124193 nm. 1D scattering intensities of samples and buffers were expressed as a function of the modulus of the scattering vector *Q* = (4π/λ)sinθ with 2θ being the scattering angle and λ the X-ray wavelength. Downstream processing after buffer subtraction was done with PRIMUS^90^. *R*g was determined using Guinier approximation and from p(*r*) curve. Disordered regions in the crystal structure were modelled using CORAL^51^ and subsequently crystal structure validation was done using CRYSOL^91^.

### Isothermal titration calorimetry

ITC experiments were performed with MicroCal PEAQ-ITC calorimeter (Malvern). Protein samples were dialyzed overnight in buffer containing 20 mM HEPES pH 7.5, 150 mM NaCl, 2 mM BME. The cell was filled completely with 40-50 µM protein and the syringe was filled with concentrations in the range of 400-500 µM of the respective ligand. A series of 19 or 26 injections of 2 µl or 1.5 µl titrants were made into the respective protein. For competition experiments, Pla1CTD was pre-incubated with a 1.1-fold molar excess of either Red1_288-345_ or Iss147-70 before loading the sample into the cell. The data were processed with PEAQ-ITC Analysis software (PEAQ-ITC) and were fit to a one-binding site model.

### *In vitro* polyadenylation assay

A 15 nt polyA chemically synthesized RNA primer with a 5’ Cy3 fluorophore (purchased from iba Life Sciences) was used as a substrate for Pla1. Polyadenylation assays were performed in buffer containing 10 mM HEPES pH 7.5, 20 mM KCl, 5 mM MgCl2, 0.01 mM EDTA, 5% glycerol and 1mM DTT at 30 °C. 75 nM Pla1 alone or in complex with Red1_288-345_ or Red1_W298A/F305A_ at a ∼10-fold molar excess was incubated with 200 nM RNA primer for 5 min at 30 °C before starting the reaction with the addition of 2mM ATP. 10 µl fractions were collected at different time points and mixed with 10 µl denaturing formamide loading dye to stop the reaction. Products were analyzed on 14% denaturing urea-PAGE, run in 0.5x Tris-borate-EDTA buffer for 45 min at 35 mA and the gels were imaged using Amersham Imager 600 (GE Healthcare). Densitometric analyses for integration of gel lane intensity was performed using Fiji ImageJ software.

### Yeast two hybrid

The respective sequences of the proteins were cloned into pGBKT7 and pGADT7 vectors from Clontech which harbor either the DNA binding domain or activation domain at the N-termini, respectively. The interaction pairs were analyzed by co-transformation into PJ69-4A strain. After a 10-fold serial dilution, colonies were spotted on SDC (SDC-Leu-Trp), SDC-His (SDC-Leu-Trp-His) and SDC-Ade (SDC-Leu-Trp-Ade) plates, incubated at 30 °C and analyzed after 3 days. The strength of the interaction was assessed by growth achieved on SDC-His and SDC-Ade as weak and strong, respectively.

### Bio-layer Interferometry

BLI experiments were performed on Octet RED96e system (Fortébio) at 25 °C in buffer containing 20 mM HEPES pH 7.5, 150 mM NaCl, 2 mM BME and 0.01% Tween-20. The protein ligands (Red1_288-345_ and Iss1) were biotinylated using EZ-Link NHS-PEG4-Biotin (Thermo Fisher Scientific). Biotin-labeled proteins were immobilized on the streptavidin (SA) biosensors (Fortébio) and a 2-fold serial dilution of the respective analyte was applied to the biosensors. Parallel experiments with a reference sensor where no analyte was added served as control and its signal was subtracted during data analysis. The association and dissociation periods were both set to 240 sec. Data measurements and analysis were performed by using the Data analysis HT 10.0 (Fortébio) software, with a global (full) 1:1 fitting model.

For studies of Pla1-A15 RNA interactions, a 15 nt RNA ligand (A15) was purchased with a 5’-BiotinTEG modification from Integrated DNA Technologies (IDT) and immobilized on the SA biosensors. Experiments were performed in buffer containing 20 mM HEPES pH 7.5, 150 mM NaCl, 2 mM BME, 0.01% Tween-20 and 0.1% Bovine Serum Albumin and the association and dissociation periods were both set to 100 sec. Experiments in the presence of Red1 were performed by adding 1.2 µM Red1_288-345_ directly in the assay buffer.

### Tandem affinity purification followed by interaction analysis/ MS

Flag-TEV-protein A (FTP)-tagged bait proteins were harvested from 2 L YEA cultures of yeast strains grown to OD_600_ 1.8–2.2. Cell pellets were snap-frozen in liquid nitrogen and ground into powder using the Cryo-mill MM-400 (Retsch machine). Cells were resuspended in purification buffer (50 mM HEPES, pH 7.0; 100 mM NaCl; 1.5 mM MgCl2; 0.15% NP-40), supplemented with 1 mM dithiothreitol (Sigma), 1 mM phenylmethylsulphonyl fluoride (Sigma Aldrich, 78830) and protease inhibitor mix (SERVA, 39104). Cell extracts were centrifuged at 3500 g for 10 min at 4 °C, then the supernatants were further centrifuged at 27000 g for 45 min at 4 °C. Clarified supernatants were then incubated with 150 mL slurry of IgG beads (GE Healthcare, 17-0969-02) for 2h at 4 °C on a turning wheel. After binding, the beads were washed with 2 x 15 mL purification buffer and TEV cleavage was performed in purification buffer containing 20 units AcTEV protease (Invitrogen 12575015), 0.5 mM dithiothreitol and 50 units Benzonase (Sigma, E1014) for the removal of nucleic acids, for 2 h at 16 °C. The eluate was collected and incubated with 100 mL slurry of anti-Flag beads (Sigma Aldrich, A2220) for 1h at 4 °C. The protein-bound anti-Flag beads were washed with 2 x 10 mL purification buffer and the proteins were subsequently eluted from the beads by competition with 200 mL Flag peptide (Assay Matrix, A6001). The resulting eluate was collected and used for total protein isolation using Trichloroacetic acid-TCA (Sigma, T0699). The eluted proteins were then analysed by Coomassie staining using Brilliant Blue G colloidal concentrate (Sigma Aldrich, B2025) on 4–12% NuPAGE Bis-Tris gels (Life Technologies).

For interaction analysis, the protein samples separated by SDS-PAGE gel was transferred onto a nitrocellulose (Invitrogen, IB301002) membrane and was probed with respective antibodies. For mass spectrometry analysis, the precipitated protein samples were subjected to an in-solution tryptic digest using a modified version of the Single-Pot Solid-Phase-enhanced Sample Preparation (SP3) protocol^92, 93^. In total three biological replicates were prepared including control, wild-type and mutant derived lysates (n=3). Lysates were added to Sera-Mag Beads (Thermo Scientific, #4515-2105-050250, 6515-2105-050250) in 10 µl 15% formic acid and 30 µl of ethanol. Binding of proteins was achieved by shaking for 15 min at room temperature (RT). SDS was removed by 4 subsequent washes with 200 µl of 70% ethanol. Proteins were digested overnight at room temperature with 0.4 µg of sequencing grade modified trypsin (Promega, #V5111) in 40 µl Hepes/NaOH, pH 8.4 in the presence of 1.25 mM TCEP and 5 mM chloroacetamide (Sigma-Aldrich, #C0267). Beads were separated, washed with 10 µl of an aqueous solution of 2% DMSO and the combined eluates were dried down. Peptides were reconstituted in 10 µl of H_2_O and reacted for 1 h at room temperature with 80 µg of TMT10plex (Thermo Scientific, #90111)^94^ label reagent dissolved in 4 µl of acetonitrile. Excess TMT reagent was quenched by the addition of 4 µl of an aqueous 5% hydroxylamine solution (Sigma, 438227). Peptides were reconstituted in 0.1 % formic acid, mixed to achieve a 1:1 ratio across all TMT-channels and purified by a reverse phase clean-up step (OASIS HLB 96-well µElution Plate, Waters #186001828BA). Peptides were subjected to an off-line fractionation under high pH conditions^93^. The resulting 12 fractions were then analyzed by LC-MS/MS on an Orbitrap Fusion Lumos mass spectrometer (Thermo Scentific) as previously described^95^. To this end, peptides were separated using an Ultimate 3000 nano RSLC system (Dionex) equipped with a trapping cartridge (Precolumn C18 PepMap100, 5 mm, 300 μm i.d., 5 μm, 100 Å) and an analytical column (Acclaim PepMap 100. 75 × 50 cm C18, 3 mm, 100 Å) connected to a nanospray-Flex ion source. The peptides were loaded onto the trap column at 30 µl per min using solvent A (0.1% formic acid) and eluted using a gradient from 2 to 40% Solvent B (0.1% formic acid in acetonitrile) over 2 h at 0.3 µl per min (all solvents were of LC-MS grade). The Orbitrap Fusion Lumos was operated in positive ion mode with a spray voltage of 2.4 kV and capillary temperature of 275 °C. Full scan MS spectra with a mass range of 375–1500 m/z were acquired in profile mode using a resolution of 120,000 (maximum fill time of 50 ms or a maximum of 4e5 ions (AGC) and a RF lens setting of 30%. Fragmentation was triggered for 3 s cycle time for peptide like features with charge states of 2–7 on the MS scan (data-dependent acquisition). Precursors were isolated using the quadrupole with a window of 0.7 m/z and fragmented with a normalized collision energy of 38. Fragment mass spectra were acquired in profile mode and a resolution of 30,000 in profile mode. Maximum fill time was set to 64 ms or an AGC target of 1e5 ions). The dynamic exclusion was set to 45 s.

Acquired data were analyzed using IsobarQuant^96^ and Mascot V2.4 (Matrix Science) using a reverse UniProt FASTA Schizosaccharomyces pombe database (UP000002485) including common contaminants. The following modifications were taken into account: Carbamidomethyl (C, fixed), TMT10plex (K, fixed), Acetyl (N-term, variable), Oxidation (M, variable) and TMT10plex (N-term, variable). The mass error tolerance for full scan MS spectra was set to 10 ppm and for MS/MS spectra to 0.02 Da. A maximum of 2 missed cleavages were allowed. A minimum of 2 unique peptides with a peptide length of at least seven amino acids and a false discovery rate below 0.01 were required on the peptide and protein level^97^, resulting in 164 proteins. Raw TMT reporter ion intensities (signal_sum columns) were first cleaned for batch effects using limma^98^ and further normalized using vsn (variance stabilization normalization^99^. Proteins were tested for differential expression using the limma package. The replicate information was not added as a factor in the design matrix since some condition were measured with a single replicate only. A protein was annotated as a hit with a false discovery rate (fdr) smaller 5 % and a fold-change of at least 100 % and as a candidate with a fdr below 20 % and a fold-change of at least 50 %.

### Total RNA isolation

Total RNAs were isolated using TriReagent (Sigma-Aldrich, T9424). Briefly, cell pellets measuring an OD_600_ of 8 was resuspended in Tri Reagent and lysed, treated twice using 1-Bromo-3-chloropropane (Sigma-Aldrich, B9673) followed by RNA precipitation using 2-propanol (Sigma-Aldrich). The RNA pellet was washed in ice-cold 75% ethanol and solubilized in nuclease-free water. The RNA concentration was measured using NanoDrop (Thermo Scientific) and 10μg of RNA was selected for DNase treatment using NEB DNase 1 (M0303). The DNase treated samples were purified further using the RNA Clean and Concentrator Kit (Zymo Research, R1015/R1017) and stored at – 80°C after addition of ribonuclease inhibitor (Invitrogen, 10777019).

### Illumina sequencing of poly(A)+ RNA

Total RNA was isolated, DNase treated and purified from respective strains as mentioned above. The RNA quality was assessed using Bioanalyzer (Agilent 2100) and only those of high quality was proceeded to library preparation. Briefly, 1 μg was taken from individual samples, and ERCC-RNA spikeIn (Life technologies, 4456740) was added according to manufacturer’s instructions. The RNA samples were then subjected to OligodT purification of poly(A) RNA (NEB, E7490) following the manufacturer’s instructions and the recovered poly(A) selected RNA was used for cDNA library preparation using NEBNext Ultra II directional RNA library kit (NEB, E7760) following library preparation protocol. The final cDNA libraries were amplified (cycle number of 8) using NEBNext Multiplex Oligos for Illumina (NEB, E7335) and purified using SPRIselect size selection beads (Beckman Coulter, B23317). Individual library quality was assessed on TapeStation (Agilent 4200) using a DNA D1000 High sensitivity tape. Indexed libraries were pooled and sequenced with ∼20M reads per sample on a NovaSeq. sequencer with 150 bp paired-end reads.

### Illumina sequencing of total RNA

Total RNA was isolated, DNase treated and purified from respective strains as mentioned above. The RNA quality was assessed using Bioanalyzer (Agilent 2100) and only those of high quality was proceeded to library preparation. Briefly, 100ng was taken from individual samples, and an ERCC-RNA spikeIn (Life technologies, 4456740) amount equivalent to that for 1μg of RNA input (equivalent to that of poly(A)+ sample) was added. The RNA samples were directly proceeded for cDNA library preparation using NEBNext Ultra II directional RNA library kit (NEB, E7760) following the library preparation protocol. The final cDNA libraries were amplified (cycle number of 6) using NEBNext Multiplex Oligos for Illumina (NEB, E7335) and purified using SPRIselect size selection beads (Beckman Coulter, B23317). Individual library quality was assessed on TapeStation (Agilent 4200) using a DNA D1000 High sensitivity tape. Indexed libraries were pooled and subjected for deep sequencing with ∼80M reads per sample on a NovaSeq. sequencer with 150 bp paired-end reads.

### Nanopore sequencing of poly(A)+ RNA

Total RNA was isolated, DNase treated and purified from respective strains as mentioned above. The RNA quality was assessed using Bioanalyzer (Agilent 2100) and only those of high quality was proceeded to library preparation. Briefly, 5μg of RNA samples was subjected to OligodT purification of poly(A) RNA (NEB, E7490) following the manufacturer’s instructions and 100ng of recovered poly(A)+ selected RNA was used for library preparation using Oxford Nanopore direct RNA sequencing library kit (SQK-RNA002; Version: DRS_9080_v2_revM_14Ag2019) protocol. The prepared libraries were run on individual flow cells following manufacturer guidelines.

### RNA-Immunoprecipitation (RIP) followed by sequencing (Illumina and Nanopore)

Tandem affinity purifications of FTP-tagged strains were performed as described above with minor modifications. Reagents were prepared under RNase-free conditions, in the presence of RNase inhibitor (Invitrogen, 10777019). Two-third of final flag elute was taken for DNase treatment using NEB DNase 1 (M0303) and RNA was purified using RNA Clean and Concentrator Kit (Zymo Research, R1015/R1017). The final RNA concentration was determined using Qubit RNA high Sensitivity Assay Kit and 100ng of RIP-purified RNA was used for cDNA library preparation using NEBNext Ultra II directional RNA library kit (NEB, E7760) following library preparation protocol. The final cDNA libraries were amplified using NEBNext Multiplex Oligos for Illumina (NEB, E7335) and purified using SPRIselect size selection beads (Beckman Coulter, B23317). Individual library quality was assessed on TapeStation (Agilent 4200) using a DNA D1000 High sensitivity tape. Indexed libraries were pooled and sequenced on NovaSeq sequencer with 150 bp paired-end reads. Similarly, 200ng of RIP-purified RNA was used for library preparation using Oxford Nanopore direct RNA sequencing library kit (SQK-RNA002; Version: DRS_9080_v2_revM_14Ag2019) protocol. The prepared libraries were run on individual flow cells following manufacturer guidelines.

### RNA-seq analysis

Paired-end illumina reads were aligned with hisat 2.1.0^100^ allowing introns with a maximum length of 2,000 nt (‘--max.intronlen 2000’). Aligned bam files were then sorted and indexed using samtools 1.3.1 ^101^ Nanopore reads were aligned with minimap (2.10) with long-read spiced alignment (with splice and -k7 parameters). Strand-specific bigwig tracks were generated using bamCoverage ^102^ to the pombe genome (ASM294v2) with the parameter ‘—binSize 1’. Bigwig tracks were normalised by the sum of the raw signals of chromosome I and II multiplied by 100 million. Meta plots were generated with in-house R scripts using GenomicRanges^103^ packages. Integrative Genomics Viewer (IGV2.3, Broad Institute) was used for data browsing (32) and creating representative snapshots.

### Detection and quantification of CUTs and PROMPTs

Putative CUT regions were created by neighbouring signals closer than 25nts and showing FC>1.2 relative to red11β (length>=100 nts) using Rsamtools ^104^ and dplyr ^105^ separately on the forward and reverse bigwig tracks. PROMPT regions were defined as CUTs that are shorter than 1500 nts and in the +-250nts vicinity of an annotated pombe gene (ASM294v2) on the opposite strand. Intersections, subtractions and merging of the predicted regions were done with BedTools 2.28.0.

### Quantification of polyA length from poly(A)+ RNA Nanopore sequencing

polyA length for the whole transcriptome, meiotic genes, CUT and PROMPT regions was estimated using tailfindr^54^ and boxplots of arithmetic mean of signals per regions were plotted with the ggplot2 R package.

### Chromatin Immunoprecipitation-sequencing (ChIP-seq.)

*S. pombe* strains grown to an OD_600_ of 0.5-0.8 were crosslinked using formaldehyde solution (Sigma, F1635) to a final concentration of 1% at RT for 15 min and quenched by addition of glycine (ThermoFisher, AJA1083) to a final concentration of 150 mM for 5 min at RT. The crosslinked culture was then washed and harvested twice using 1x PBS (137mM NaCl, 2.7mM KCl, 10mM Na_2_HPO_4_, 1.2mM KH_2_PO_4_) by centrifugation at 2600 rpm for 2 min at 4°C each and the final cell pellet was frozen in liquid nitrogen. The cell pellet was then resuspended in 300 μL FA-SDS buffer (50 mM HEPES-KOH pH 8, 2 mM EDTA pH 8, 150 mM NaCl, 1% TritonX-100, 0.1% Sodium Deoxycholate, 0.1% SDS) supplemented with 1x protease inhibitor mix (SERVA, 39104), 1 mM phenylmethylsulfonyl fluoride (Sigma Aldrich, 78830) was homogenized at 4°C using 700 µL zirconia beads (BioSpec, 110791) in Precellys 24 homogenizer (Bertin Technologies, France) at 5500 rpm. The lysate was collected and transferred into the Covaris AFA Fiber&cap 12×12mm millitube (Covaris, 520135) after bringing the final amount to 1 mL total. The chromatin was then sheared to a median size of ∼300bp by sonication using Covaris S2 (Adaptive Focused Acoustics™ (AFA) technology) at the given parameters: Duty cycle: 20%, Intensity: 5, Cycles/burst: 200, Temp: 7°C, Time: 6 min. The sonicated lysate was collected, centrifuged at 3000 rpm for 10 minutes at 4°C and the supernatant collected was diluted at 1:1 ratio using FA-lysis buffer (50 mM HEPES-KOH pH 7.5, 2 mM EDTA pH 7.5, 150 mM NaCl, 1% TritonX-100, 0.1% Sodium Deoxycholate) to dilute the SDS concentration to 0.05%. The samples were then incubated overnight at 4°C with 2μ<u>g</u> of H3K9me2 antibody (mAbcam 1220).

Following antibody binding, the samples were incubated at 4°C for at least 1 hr with ∼40 µL of appropriate bead slurry (nProtein A Sepharose, GE Healthcare, 5280-01). Following incubation, the beads were collected by centrifugation at 4°C at 1200 rpm for 1 min. The beads were then washed twice with washing buffer I (50 mM HEPES-KOH pH 7.5, 1 mM EDTA pH 7.5, 140 mM NaCl, 1% TritonX-100, 0.1% Sodium Deoxycholate) and wash buffer II (50 mM HEPES-KOH pH 7.5, 1 mM EDTA pH 7.5, 0.5 M NaCl, 1% TritonX-100, 0.1% Sodium Deoxycholate). Finally, the beads were washed once again with wash buffer III (10 mM Tris-Cl pH 8, 1 mM EDTA pH 7.5, 250 mM LiCl, 1% NP-40 Igepal, 0.5% Sodium Deoxycholate) and briefly with 1 mL Tris-EDTA pH 8 buffer. The bound protein and chromatin fractions were eluted twice by adding 50 µL of elution buffer (10 mM Tris-HCl pH 8, 10 mM EDTA pH 7.5, 2% SDS) at 65°C for 10-15 min at 1000 rpm on a thermomixer. The samples were then de-crosslinked by incubating at 65°C overnight after addition of 2 µL proteinase K. The DNA was purified using Phenol: Chloroform: Isomyl alcohol method following RNase treatment. The purified DNA pellet was eluted in 15-20 µL of nuclease free water (Invitrogen, 10977015) or 0.1x TE and was used for library preparation using NEBNext Ultra II DNA library prep kit for Illumina (NEB, E7645) following manufacturer instructions. The final cDNA libraries were amplified using NEBNext Multiplex Oligos for illumina (NEB, E7335) and purified using SPRIselect size selection beads (Beckman Coulter, B23317). Individual library quality was assessed on TapeStation (Agilent 4200) using a DNA D1000 High sensitivity tape. Indexed libraries were pooled and sequenced on NovaSeq sequencer with 150 bp paired-end reads.

### ChIP-seq analysis

Paired-end illumina reads were subject to a thorough Quality Control (QC). Shortly, FastQC 0.11.8 ^106^ as used to generate QC reports. Low complexity regions were filtered out with prinseq-lite.pl from PRINSEQ-lite 0.20.4^107^ maximum dust was set to 3) and the remaining sequences were trimmed using Trimmomatic^108^ (SLIDINGWINDOW: 4 nt; phred quality cut-off: 15, MINLEN=36). QC filtered were aligned with BWA MEM -0.7.17^109^. Aligned bam files were sorted and indexed using samtools 1.3.1^101^ Potential PCR duplicates were removed with samtools using the rmdup option. Bigwig tracks were generated using bamCoverage ^102^ to the pombe genome (ASM294v2) with the parameter “—binSize 1”. Bigwig tracks were normalised by the sum of the raw signals of chromosome I and II multiplied by 100 million. Potential artifacts were defined as regions consisting of less than 15 edges and subsequently removed with a custom R script using rtracklayer^110^ and GenomicRanges^103^ packages. Meta plots were generated using in-house R-based pipeline.

### Generating the Pla1 interaction mutants

The *Red1_W298A/F305A_* and *Red11__Δ288-345__* mutants were generated using conventional primer-based cloning. Together with the wild-type Red1, these constructs were cloned into a pFa6a-Hygro plasmid in untagged and 3xFTP tagged versions. The primers used to generate the mutations are listed in Supplementary Table S4. The plasmids generated were confirmed for mutations using PCR and Sanger sequencing at the ACRF Biomolecular Resource Facility at JCSMR, ANU. The desired cassettes were then digested from the positive plasmid using Spe1 restriction enzyme that cuts in the 5’UTR and 3’UTR sequence generating overlapping region for homologous recombination in the Red1 locus when used to transform with a red11β strain.

In-order to generate strains retaining the 3’UTR, 300bp starting from the stop codon of Red1 gene was amplified along with a 500bp homology region upstream. Similarly, 500bp region following the 300bp was amplified and Gibson assembled to PCR amplified product from pFA6a-NatNT2 using appropriate primer pairs. This overrode the Hygromycin resistance marker to a natNT2 resistance along with retaining 300bp 3’UTR region of Red1. Hence all the untagged mutants used for RNA expression analysis were constructed retaining their 3’UTR region.

## Data availability

NMR backbone chemical shifts for Red1 have been deposited to the BMRB under the accession code 50680. Coordinates and structure factors for the Pla1-Red1 complex, Pla1_FL_ and Pla11β14 have been deposited in the PDB with accession codes 7Q72, 7Q73 and 7Q74, respectively. SAXS data for Pla1-Red1 complex have been deposited to the SASBDB with accession code SASDKE6. Genome-wide datasets are deposited in NCBI GEO under the reference number GEO: GSE206106 with reviewer access token: qnatoqqyxtubnqn

## Code availability

Custom R-based codes that were used to analyse the data will be deposited in Zenodo, and in the meantime are available for reviewers upon request.

## Author contributions

K.S., A.S., T.F. and I.S. designed the study and interpreted the results. K.S. performed NMR, crystallographic structure determination, SAXS, *in vitro* polyadenylation assay, biophysical characterization using pull-down assays, ITC, BLI and Y2H. N.D. and L.K. performed Y2H and NMR experiments. A.S. performed TAP purification followed by interaction analysis and mass spectrometry, Illumina sequencing of total, poly(A)+ and RIP-ed RNA, Direct RNA sequencing by Nanopore of poly(A)+ and RIP-ed RNA, ChIP seq. experiments. R.H. supported Nanopore sequencing experiments. A.H. and T.F. performed the bioinformatics analyses. B.S. recorded NMR experiments. K.W. supported crystallographic analysis. K.S, I.S and T.F wrote the manuscript. All authors contributed to the final version of the manuscript.

## Acknowledgements

We thank Jürgen Kopp and Claudia Siegmann from the BZH/Cluster of Excellence: CellNetworks crystallization platform and acknowledge access to the beamlines P12, P13 and P14 at PETRA III at DESY in Hamburg and the support of the beamline scientists. We thank Per Haberkant and Frank Stein at the Proteomics Core Facility (PCF), EMBL Heidelberg for mass spectrometry analysis. This work was supported by a DAAD fellowship to N.D., the Deutsche Forschungsgemeinschaft (DFG) through the Leibniz program (SI 586/6-1) and TRR 319 to I.S. and the Australian Research Council’s Discovery Projects funding scheme to TF (project DP190100423).

## References

1. Kilchert, C., Wittmann, S. & Vasiljeva, L. The regulation and functions of the nuclear RNA exosome complex. Nat Rev Mol Cell Biol 17, 227–39 (2016).

2. Mitchell, P. Exosome substrate targeting: the long and short of it. Biochem Soc Trans 42, 1129–34 (2014).

3. Mitchell, P., Petfalski, E., Shevchenko, A., Mann, M. & Tollervey, D. The exosome: a conserved eukaryotic RNA processing complex containing multiple 3’-->5’ exoribonucleases. Cell 91, 457–66 (1997).

4. Core, L.J., Waterfall, J.J. & Lis, J.T. Nascent RNA sequencing reveals widespread pausing and divergent initiation at human promoters. Science 322, 1845–8 (2008).

5. Xu, Z. et al. Bidirectional promoters generate pervasive transcription in yeast. Nature 457, 1033–7 (2009).

6. Neil, H. et al. Widespread bidirectional promoters are the major source of cryptic transcripts in yeast. Nature 457, 1038–42 (2009).

7. Porrua, O. & Libri, D. Transcription termination and the control of the transcriptome: why, where and how to stop. Nat Rev Mol Cell Biol 16, 190–202 (2015).

8. Davis, C.A. & Ares, M., Jr. Accumulation of unstable promoter-associated transcripts upon loss of the nuclear exosome subunit Rrp6p in Saccharomyces cerevisiae. Proc Natl Acad Sci U S A 103, 3262–7 (2006).

9. Houalla, R. et al. Microarray detection of novel nuclear RNA substrates for the exosome. Yeast 23, 439–54 (2006).

10. Zinder, J.C. & Lima, C.D. Targeting RNA for processing or destruction by the eukaryotic RNA exosome and its cofactors. Genes Dev 31, 88–100 (2017).

11. Schneider, C. & Tollervey, D. Threading the barrel of the RNA exosome. Trends Biochem Sci 38, 485–93 (2013).

12. Jia, H. et al. The RNA helicase Mtr4p modulates polyadenylation in the TRAMP complex. Cell 145, 890–901 (2011).

13. Wlotzka, W., Kudla, G., Granneman, S. & Tollervey, D. The nuclear RNA polymerase II surveillance system targets polymerase III transcripts. EMBO J 30, 1790–803 (2011).

14. Jackson, R.N. et al. The crystal structure of Mtr4 reveals a novel arch domain required for rRNA processing. EMBO J 29, 2205–16 (2010).

15. Callahan, K.P. & Butler, J.S. TRAMP complex enhances RNA degradation by the nuclear exosome component Rrp6. J Biol Chem 285, 3540–7 (2010).

16. Ogami, K., Chen, Y. & Manley, J.L. RNA surveillance by the nuclear RNA exosome: mechanisms and significance. Noncoding RNA 4(2018).

17. LaCava, J. et al. RNA degradation by the exosome is promoted by a nuclear polyadenylation complex. Cell 121, 713–24 (2005).

18. Wyers, F. et al. Cryptic pol II transcripts are degraded by a nuclear quality control pathway involving a new poly(A) polymerase. Cell 121, 725–37 (2005).

19. Mohanty, B.K. & Kushner, S.R. Bacterial/archaeal/organellar polyadenylation. Wiley Interdiscip Rev RNA 2, 256–76 (2011).

20. Eckmann, C.R., Rammelt, C. & Wahle, E. Control of poly(A) tail length. Wiley Interdiscip Rev RNA 2, 348–61 (2011).

21. Anderson, J.T. & Wang, X. Nuclear RNA surveillance: no sign of substrates tailing off. Crit Rev Biochem Mol Biol 44, 16–24 (2009).

22. Tudek, A., Lloret-Llinares, M. & Jensen, T.H. The multitasking polyA tail: nuclear RNA maturation, degradation and export. Philos Trans R Soc Lond B Biol Sci 373(2018).

23. Lubas, M. et al. Interaction profiling identifies the human nuclear exosome targeting complex. Mol Cell 43, 624–37 (2011).

24. Meola, N. et al. Identification of a Nuclear Exosome Decay Pathway for Processed Transcripts. Mol Cell 64, 520–533 (2016).

25. Lee, N.N. et al. Mtr4-like protein coordinates nuclear RNA processing for heterochromatin assembly and for telomere maintenance. Cell 155, 1061–74 (2013).

26. Zhou, Y. et al. The fission yeast MTREC complex targets CUTs and unspliced pre-mRNAs to the nuclear exosome. Nat Commun 6, 7050 (2015).

27. Egan, E.D., Braun, C.R., Gygi, S.P. & Moazed, D. Post-transcriptional regulation of meiotic genes by a nuclear RNA silencing complex. RNA 20, 867–81 (2014).

28. Silla, T., Karadoulama, E., Makosa, D., Lubas, M. & Jensen, T.H. The RNA Exosome Adaptor ZFC3H1 Functionally Competes with Nuclear Export Activity to Retain Target Transcripts. Cell Rep 23, 2199–2210 (2018).

29. Dobrev, N. et al. The zinc-finger protein Red1 orchestrates MTREC submodules and binds the Mtl1 helicase arch domain. Nat Commun 12, 3456 (2021).

30. Christofori, G. & Keller, W. 3’ cleavage and polyadenylation of mRNA precursors in vitro requires a poly(A) polymerase, a cleavage factor, and a snRNP. Cell 54, 875–89 (1988).

31. Takagaki, Y., Ryner, L.C. & Manley, J.L. Separation and characterization of a poly(A) polymerase and a cleavage/specificity factor required for pre-mRNA polyadenylation. Cell 52, 731–42 (1988).

32. Gilmartin, G.M. & Nevins, J.R. An ordered pathway of assembly of components required for polyadenylation site recognition and processing. Genes Dev 3, 2180–90 (1989).

33. Yamanaka, S., Yamashita, A., Harigaya, Y., Iwata, R. & Yamamoto, M. Importance of polyadenylation in the selective elimination of meiotic mRNAs in growing S. pombe cells. EMBO J 29, 2173–81 (2010).

34. Chen, H.M., Futcher, B. & Leatherwood, J. The fission yeast RNA binding protein Mmi1 regulates meiotic genes by controlling intron specific splicing and polyadenylation coupled RNA turnover. PLoS One 6, e26804 (2011).

35. Sugiyama, T. & Sugioka-Sugiyama, R. Red1 promotes the elimination of meiosis-specific mRNAs in vegetatively growing fission yeast. EMBO J 30, 1027–39 (2011).

36. Yamashita, A. et al. Hexanucleotide motifs mediate recruitment of the RNA elimination machinery to silent meiotic genes. Open Biol 2, 120014 (2012).

37. Bard, J. et al. Structure of yeast poly(A) polymerase alone and in complex with 3’-dATP. Science 289, 1346–9 (2000).

38. Martin, G., Keller, W. & Doublie, S. Crystal structure of mammalian poly(A) polymerase in complex with an analog of ATP. EMBO J 19, 4193–203 (2000).

39. Daubner, G.M., Clery, A. & Allain, F.H. RRM-RNA recognition: NMR or crystallography…and new findings. Curr Opin Struct Biol 23, 100–8 (2013).

40. Balbo, P.B. & Bohm, A. Mechanism of poly(A) polymerase: structure of the enzyme-MgATP-RNA ternary complex and kinetic analysis. Structure 15, 1117–31 (2007).

41. Balbo, P.B., Meinke, G. & Bohm, A. Kinetic studies of yeast polyA polymerase indicate an induced fit mechanism for nucleotide specificity. Biochemistry 44, 7777–86 (2005).

42. Balbo, P.B., Toth, J. & Bohm, A. X-ray crystallographic and steady state fluorescence characterization of the protein dynamics of yeast polyadenylate polymerase. J Mol Biol 366, 1401–15 (2007).

43. Kielkopf, C.L., Lucke, S. & Green, M.R. U2AF homology motifs: protein recognition in the RRM world. Genes Dev 18, 1513–26 (2004).

44. Helmling, S., Zhelkovsky, A. & Moore, C.L. Fip1 regulates the activity of Poly(A) polymerase through multiple interactions. Mol Cell Biol 21, 2026–37 (2001).

45. Preker, P.J., Lingner, J., Minvielle-Sebastia, L. & Keller, W. The FIP1 gene encodes a component of a yeast pre-mRNA polyadenylation factor that directly interacts with poly(A) polymerase. Cell 81, 379–89 (1995).

46. Forbes, K.P., Addepalli, B. & Hunt, A.G. An Arabidopsis Fip1 homolog interacts with RNA and provides conceptual links with a number of other polyadenylation factor subunits. J Biol Chem 281, 176–86 (2006).

47. Kaufmann, I., Martin, G., Friedlein, A., Langen, H. & Keller, W. Human Fip1 is a subunit of CPSF that binds to U-rich RNA elements and stimulates poly(A) polymerase. EMBO J 23, 616–26 (2004).

48. Pedersen, L.C., Benning, M.M. & Holden, H.M. Structural investigation of the antibiotic and ATP-binding sites in kanamycin nucleotidyltransferase. Biochemistry 34, 13305–11 (1995).

49. Pelletier, H., Sawaya, M.R., Kumar, A., Wilson, S.H. & Kraut, J. Structures of ternary complexes of rat DNA polymerase beta, a DNA template-primer, and ddCTP. Science 264, 1891–903 (1994).

50. Remaut, H. & Waksman, G. Protein-protein interaction through beta-strand addition. Trends Biochem Sci 31, 436–44 (2006).

51. Petoukhov, M.V. et al. New developments in the ATSAS program package for small-angle scattering data analysis. J Appl Crystallogr 45, 342–350 (2012).

52. Kuhn, U., Buschmann, J. & Wahle, E. The nuclear poly(A) binding protein of mammals, but not of fission yeast, participates in mRNA polyadenylation. RNA 23, 473–482 (2017).

53. Zhelkovsky, A., Helmling, S. & Moore, C. Processivity of the Saccharomyces cerevisiae poly(A) polymerase requires interactions at the carboxyl-terminal RNA binding domain. Mol Cell Biol 18, 5942–51 (1998).

54. Krause, M. et al. tailfindr: alignment-free poly(A) length measurement for Oxford Nanopore RNA and DNA sequencing. RNA 25, 1229–1241 (2019).

55. Lee, S.Y. et al. Dense Transposon Integration Reveals Essential Cleavage and Polyadenylation Factors Promote Heterochromatin Formation. Cell Rep 30, 2686–2698 e8 (2020).

56. Vo, T.V. et al. CPF Recruitment to Non-canonical Transcription Termination Sites Triggers Heterochromatin Assembly and Gene Silencing. Cell Rep 28, 267–281 e5 (2019).

57. Zofall, M. et al. RNA elimination machinery targeting meiotic mRNAs promotes facultative heterochromatin formation. Science 335, 96–100 (2012).

58. Yamanaka, S. et al. RNAi triggered by specialized machinery silences developmental genes and retrotransposons. Nature 493, 557–60 (2013).

59. Hiriart, E. et al. Mmi1 RNA surveillance machinery directs RNAi complex RITS to specific meiotic genes in fission yeast. EMBO J 31, 2296–308 (2012).

60. Tashiro, S., Asano, T., Kanoh, J. & Ishikawa, F. Transcription-induced chromatin association of RNA surveillance factors mediates facultative heterochromatin formation in fission yeast. Genes Cells 18, 327–39 (2013).

61. Zhao, J., Kessler, M.M. & Moore, C.L. Cleavage factor II of Saccharomyces cerevisiae contains homologues to subunits of the mammalian Cleavage/ polyadenylation specificity factor and exhibits sequence-specific, ATP-dependent interaction with precursor RNA. J Biol Chem 272, 10831–8 (1997).

62. Manley, J.L. A complex protein assembly catalyzes polyadenylation of mRNA precursors. Curr Opin Genet Dev 5, 222–8 (1995).

63. Preker, P.J., Ohnacker, M., Minvielle-Sebastia, L. & Keller, W. A multisubunit 3’ end processing factor from yeast containing poly(A) polymerase and homologues of the subunits of mammalian cleavage and polyadenylation specificity factor. EMBO J 16, 4727–37 (1997).

64. Casanal, A. et al. Architecture of eukaryotic mRNA 3’-end processing machinery. Science 358, 1056–1059 (2017).

65. Meinke, G. et al. Structure of yeast poly(A) polymerase in complex with a peptide from Fip1, an intrinsically disordered protein. Biochemistry 47, 6859–69 (2008).

66. Ezeokonkwo, C., Zhelkovsky, A., Lee, R., Bohm, A. & Moore, C.L. A flexible linker region in Fip1 is needed for efficient mRNA polyadenylation. RNA 17, 652–64 (2011).

67. Kumar, A. et al. Dynamics in Fip1 regulate eukaryotic mRNA 3’ end processing. Genes Dev (2021).

68. St-Andre, O. et al. Negative regulation of meiotic gene expression by the nuclear poly(a)-binding protein in fission yeast. J Biol Chem 285, 27859–68 (2010).

69. Makino, D.L., Baumgartner, M. & Conti, E. Crystal structure of an RNA-bound 11-subunit eukaryotic exosome complex. Nature 495, 70–5 (2013).

70. Mangus, D.A., Evans, M.C. & Jacobson, A. Poly(A)-binding proteins: multifunctional scaffolds for the post-transcriptional control of gene expression. Genome Biol 4, 223 (2003).

71. Kerwitz, Y. et al. Stimulation of poly(A) polymerase through a direct interaction with the nuclear poly(A) binding protein allosterically regulated by RNA. EMBO J 22, 3705–14 (2003).

72. Wahle, E. A novel poly(A)-binding protein acts as a specificity factor in the second phase of messenger RNA polyadenylation. Cell 66, 759–68 (1991).

73. Lemieux, C. et al. A Pre-mRNA degradation pathway that selectively targets intron-containing genes requires the nuclear poly(A)-binding protein. Mol Cell 44, 108–19 (2011).

74. Bresson, S.M. & Conrad, N.K. The human nuclear poly(a)-binding protein promotes RNA hyperadenylation and decay. PLoS Genet 9, e1003893 (2013).

75. Shi, Y. et al. Molecular architecture of the human pre-mRNA 3’ processing complex. Mol Cell 33, 365–76 (2009).

76. Tuck, A.C. & Tollervey, D. A transcriptome-wide atlas of RNP composition reveals diverse classes of mRNAs and lncRNAs. Cell 154, 996–1009 (2013).

77. Kuehner, J.N. & Brow, D.A. Regulation of a eukaryotic gene by GTP-dependent start site selection and transcription attenuation. Mol Cell 31, 201–11 (2008).

78. Gonzalez, C.I., Ruiz-Echevarria, M.J., Vasudevan, S., Henry, M.F. & Peltz, S.W. The yeast hnRNP-like protein Hrp1/Nab4 marks a transcript for nonsense-mediated mRNA decay. Mol Cell 5, 489–99 (2000).

79. Kessler, M.M. et al. Hrp1, a sequence-specific RNA-binding protein that shuttles between the nucleus and the cytoplasm, is required for mRNA 3’-end formation in yeast. Genes Dev 11, 2545–56 (1997).

80. Touat-Todeschini, L. et al. Selective termination of lncRNA transcription promotes heterochromatin silencing and cell differentiation. EMBO J 36, 2626–2641 (2017).

81. Harigaya, Y. et al. Selective elimination of messenger RNA prevents an incidence of untimely meiosis. Nature 442, 45–50 (2006).

82. Studier, F.W. Protein production by auto-induction in high density shaking cultures. Protein Expr Purif 41, 207–34 (2005).

83. Kabsch, W. Xds. Acta Crystallogr D Biol Crystallogr 66, 125–32 (2010).

84. Emsley, P., Lohkamp, B., Scott, W.G. & Cowtan, K. Features and development of Coot. Acta Crystallogr D Biol Crystallogr 66, 486–501 (2010).

85. Afonine, P.V. et al. Towards automated crystallographic structure refinement with phenix.refine. Acta Crystallogr D Biol Crystallogr 68, 352–67 (2012).

86. Tickle, I.J.F., C.; Keller, P.; Paciorek, W.; Sharff, A.; Vonrhein, C.; Bricogne, G. STARANISO (http://staraniso.globalphasing.org/cgi-bin/staraniso.cgi). Cambridge, United Kingdom: Global Phasing Ltd. (2018).

87. Sattler, M., Schleucher, J. & Griesinger, C. Heteronuclear multidimensional NMR experiments for the structure determination of proteins in solution. Progress in Nuclear Magnetic Resonance Spectroscopy 34, 93–158 (1999).

88. Delaglio, F. et al. NMRPipe: a multidimensional spectral processing system based on UNIX pipes. J Biomol NMR 6, 277–93 (1995).

89. Vranken, W.F. et al. The CCPN data model for NMR spectroscopy: development of a software pipeline. Proteins 59, 687–96 (2005).

90. Konarev, P., Volkov, V., Sokolova, A., Koch, M. & Svergun, D. PRIMUS: a Windows PC-based system for small-angle scattering data analysis. Journal of Applied Crystallography 36, 1277–1282 (2003).

91. Franke, D. et al. ATSAS 2.8: a comprehensive data analysis suite for small-angle scattering from macromolecular solutions. J Appl Crystallogr 50, 1212–1225 (2017).

92. Moggridge, S., Sorensen, P.H., Morin, G.B. & Hughes, C.S. Extending the Compatibility of the SP3 Paramagnetic Bead Processing Approach for Proteomics. J Proteome Res 17, 1730–1740 (2018).

93. Hughes, C.S. et al. Ultrasensitive proteome analysis using paramagnetic bead technology. Mol Syst Biol 10, 757 (2014).

94. Werner, T. et al. Ion coalescence of neutron encoded TMT 10-plex reporter ions. Anal Chem 86, 3594–601 (2014).

95. Sridharan, S. et al. Proteome-wide solubility and thermal stability profiling reveals distinct regulatory roles for ATP. Nat Commun 10, 1155 (2019).

96. Franken, H. et al. Thermal proteome profiling for unbiased identification of direct and indirect drug targets using multiplexed quantitative mass spectrometry. Nat Protoc 10, 1567–93 (2015).

97. Savitski, M.M., Wilhelm, M., Hahne, H., Kuster, B. & Bantscheff, M. A Scalable Approach for Protein False Discovery Rate Estimation in Large Proteomic Data Sets. Mol Cell Proteomics 14, 2394–404 (2015).

98. Ritchie, M.E. et al. limma powers differential expression analyses for RNA-sequencing and microarray studies. Nucleic Acids Res 43, e47 (2015).

99. Huber, W., von Heydebreck, A., Sultmann, H., Poustka, A. & Vingron, M. Variance stabilization applied to microarray data calibration and to the quantification of differential expression. Bioinformatics 18 **Suppl 1**, S96–104 (2002).

100. Kim, D., Langmead, B. & Salzberg, S.L. HISAT: a fast spliced aligner with low memory requirements. Nat Methods 12, 357–60 (2015).

101. Li, H. et al. The Sequence Alignment/Map format and SAMtools. Bioinformatics 25, 2078–9 (2009).

102. Ramirez, F., Dundar, F., Diehl, S., Gruning, B.A. & Manke, T. deepTools: a flexible platform for exploring deep-sequencing data. Nucleic Acids Res 42, W187–91 (2014).

103. Lawrence, M. et al. Software for computing and annotating genomic ranges. PLoS Comput Biol 9, e1003118 (2013).

104. Morgan, M.P., H.; Obenchain, V.; Hayden, N. . Rsamtools: Binary alignment (BAM), FASTA, variant call (BCF), and tabix file import. (2022).

105. Wickham, H.F., R.; Henry, L.; Müller, K. . dplyr: A Grammar of Data Manipulation. R package version. (2018).

106. Andrews, S. FastQC: a quality control tool for high throughput sequence data. (2010).

107. Schmieder, R. & Edwards, R. Quality control and preprocessing of metagenomic datasets. Bioinformatics 27, 863–4 (2011).

108. Bolger, A.M., Lohse, M. & Usadel, B. Trimmomatic: a flexible trimmer for Illumina sequence data. Bioinformatics 30, 2114–20 (2014).

109. Li, H. Aligning sequence reads, clone sequences and assembly contigs with BWA-MEM. arXiv 1303.3997v2(2013).

110. Lawrence, M., Gentleman, R. & Carey, V. rtracklayer: an R package for interfacing with genome browsers. Bioinformatics 25, 1841–2 (2009).

