## Supplementary Information for "Mechanistic insights into RNA surveillance by the canonical poly(A) polymerase Pla1 of the MTREC complex"

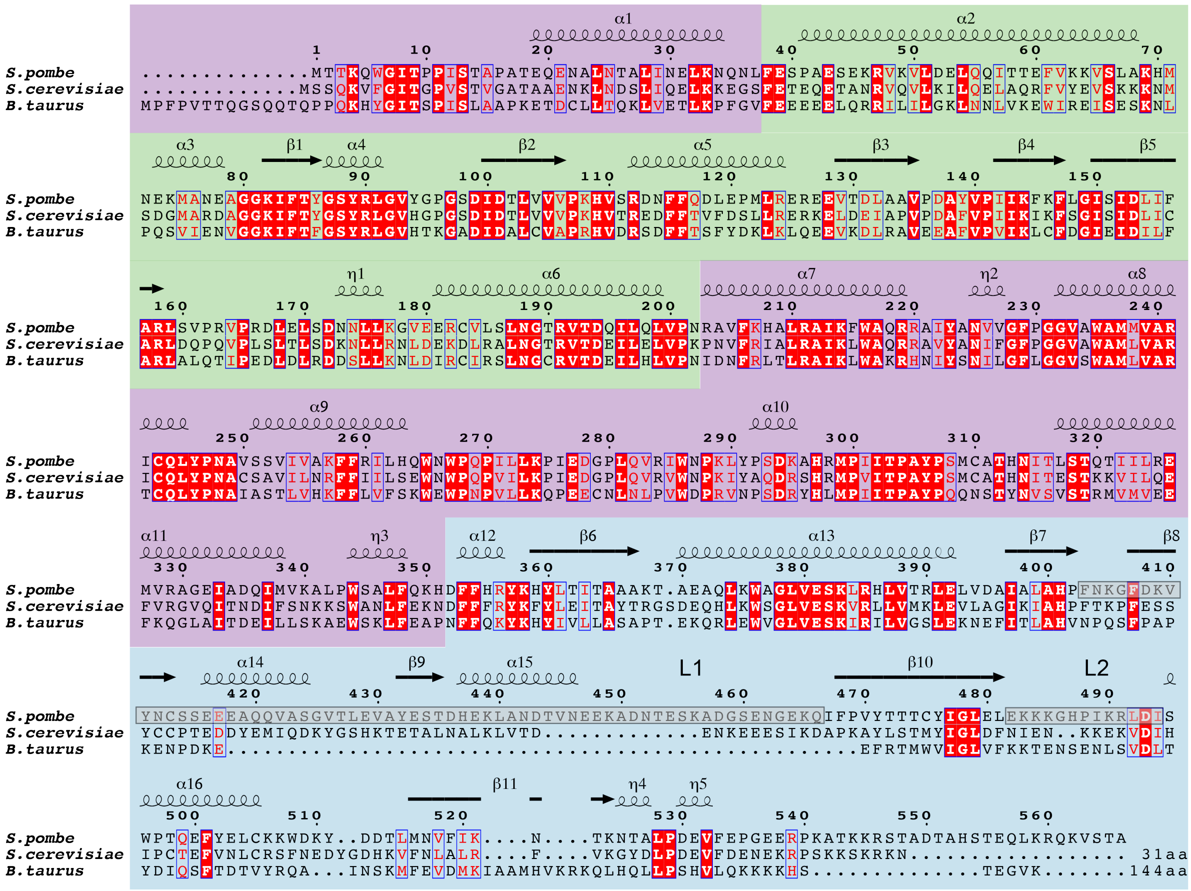

#### Supplementary Figure S1. Sequence alignment of Pla1

Sequence alignment of Pla1 from S. pombe and its homologues from *S. cerevisiae* (Pap1) and *B. taurus* (PAP) are shown. Residues which are identical are highlighted in red while those that are highly similar are colored in red and boxed in blue. 31 and 144 amino acids (aa) at the C-termini of Pap1 and PAP are not shown in the alignment. Secondary structure and sequence numbering with respect to Pla1 is shown on top of the sequence alignment. The NTD, MD and RRM domain of Pla1 are highlighted in green, purple and blue respectively. Loops L1 and L2 are marked in grey.

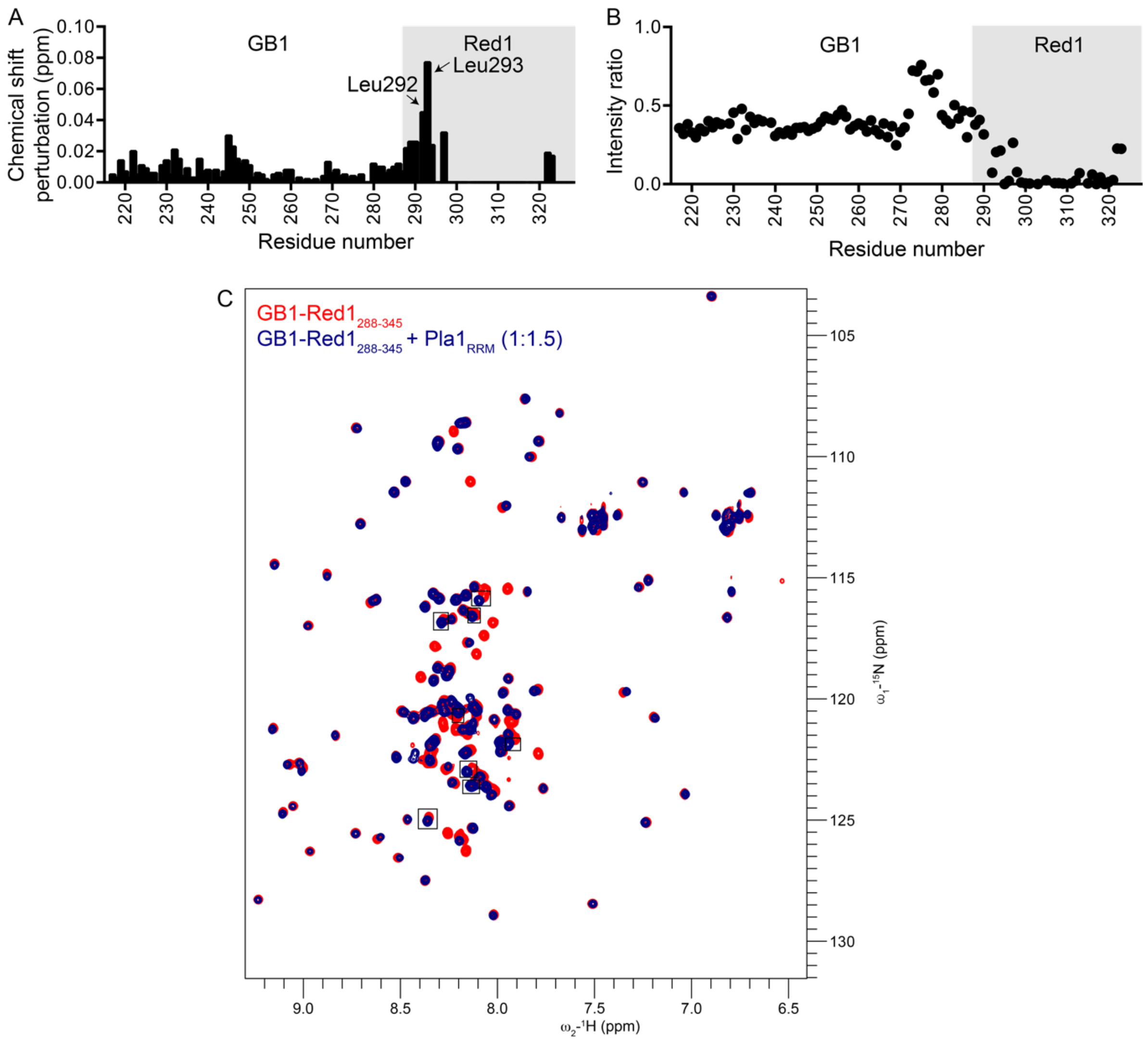


**Supplementary Figure S2. NMR analyses of Red1**

(A) Chemical shift perturbations arising in GB1-tagged Red1$_{288\text{-}322}$ in the presence of 1.5-fold molar excess of Pla1$_{RRM}$ domain. Residues belonging to Red1 are highlighted in grey. Minor CSPs are observed in the GB1 tag while more pronounced CSPs occur at the N-terminus of Red1 (residues Leu292, Leu293). (B) Intensity ratios of Red1$_{288\text{-}322}$ + Pla1$_{RRM}$ compared to free Red1$_{288\text{-}322}$ are plotted against residue number. (C) Overlay of $^{1}$H, $^{15}$N-HSQC NMR spectra of GB1-tagged Red1$_{288\text{-}345}$ in the absence (red) and presence (blue) of 1.5-fold molar excess of Pla1$_{RRM}$ domain is shown. Additional Red1 residues which show minor chemical shift perturbations in comparison to GB1-tagged Red1$_{288\text{-}322}$ (**Figure 1D**) are boxed.

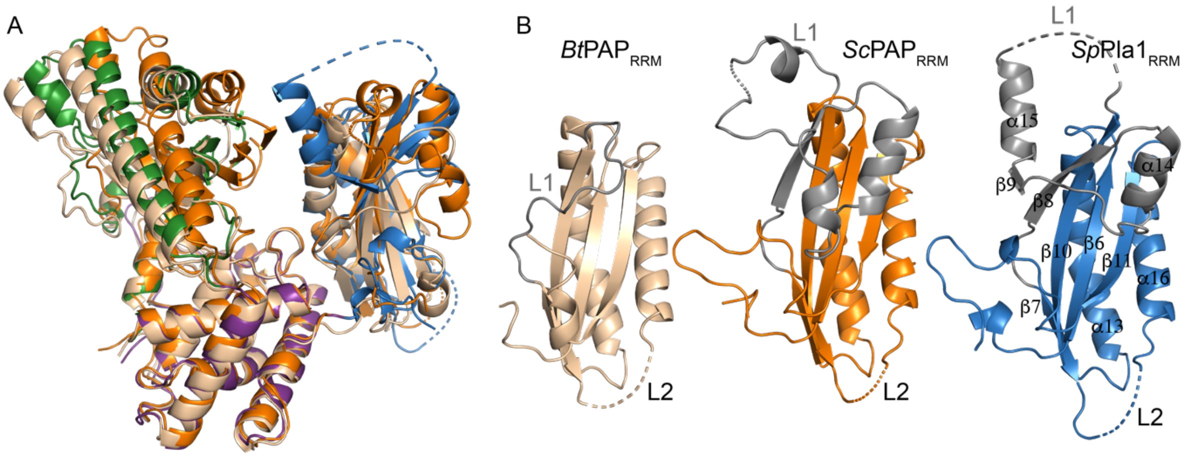


**Supplementary Figure S3. Comparison of Pla1 with its homologues**

(A) Superposition of crystal structures of Pla1 apo (color scheme based on Figure 1A) with *S. cerevisiae* Pap1 (PDB ID: 2Q66, orange) and mammalian PAP (PDB ID: 1Q79, wheat) based on an alignment on the MD. A displacement of the NTD and RRM domains shows the flexibility of the domain arrangement. (B) Similar views of the RRM domains of Pla1 and its homologues. Positions of loops L1 and L2 are marked, and L1 shown in grey. L1 is severely extended in yeast when compared to mammals, and partially blocks the canonical RNA binding β-sheet interface.

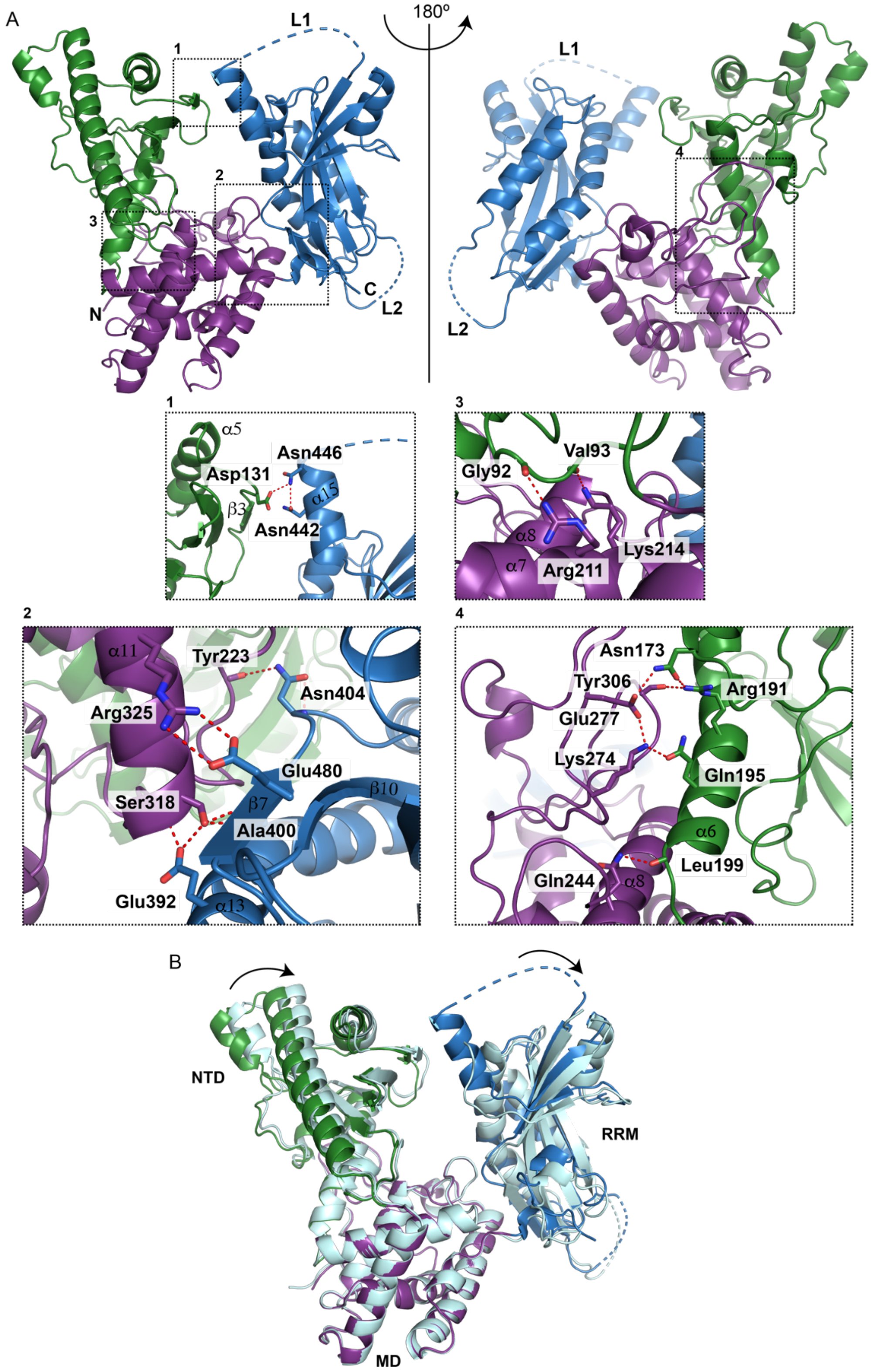


**Supplementary Figure S4. Crystal structure of Pla1**

(A). Pla1 NTD, MD and RRM domains are held together by four different interfaces which are shown in the insets. Briefly, inset 1 shows that the NTD and RRM domain are connected via salt bridges between side chains of Asp131, Asn442 and Asn446. Inset 2 shows that the MD and RRM domain are held together strongly by hydrogen bonds between side chain of Asn404 and backbone carbonyl of Tyr223, salt bridges between side chains of Glu480 and Arg325. In addition, a network of interactions between the side chain of Glu392 and backbone amide, side chain of Ser318 and Ala400 backbone is also formed. Interactions between the catalytic domain and the middle domain occur at two interfaces, as shown in insets 3 and 4. The first interface is formed by hydrogen bonds between the sidechain of Arg211 and backbone carbonyl of Gly92, side chain of Lys214 and backbone carbonyl of Val93. The second interface involves a number of interactions mediated via the helix $\alpha$6 of the NTD. Hydrogen bonds are formed between the side chains of Lys274 and Gln195, side chain of Gln244 and backbone carbonyl of Leu199, side chains of Glu277 and Asn173, backbone carbonyl of Tyr306 and side chain of Arg191. Hydrogen bonds are represented by red dotted lines. (B) Superposition of $Pla1_{FL}$ and $Pla1_{\Delta 14}$ crystal structures based on alignment of the MD. $Pla1_{FL}$ NTD, MD and RRM domain are shown in green, purple and blue, respectively while $Pla1_{\Delta 14}$ is shown in cyan. Displacement of the NTD and RRM domains is shown with arrows.

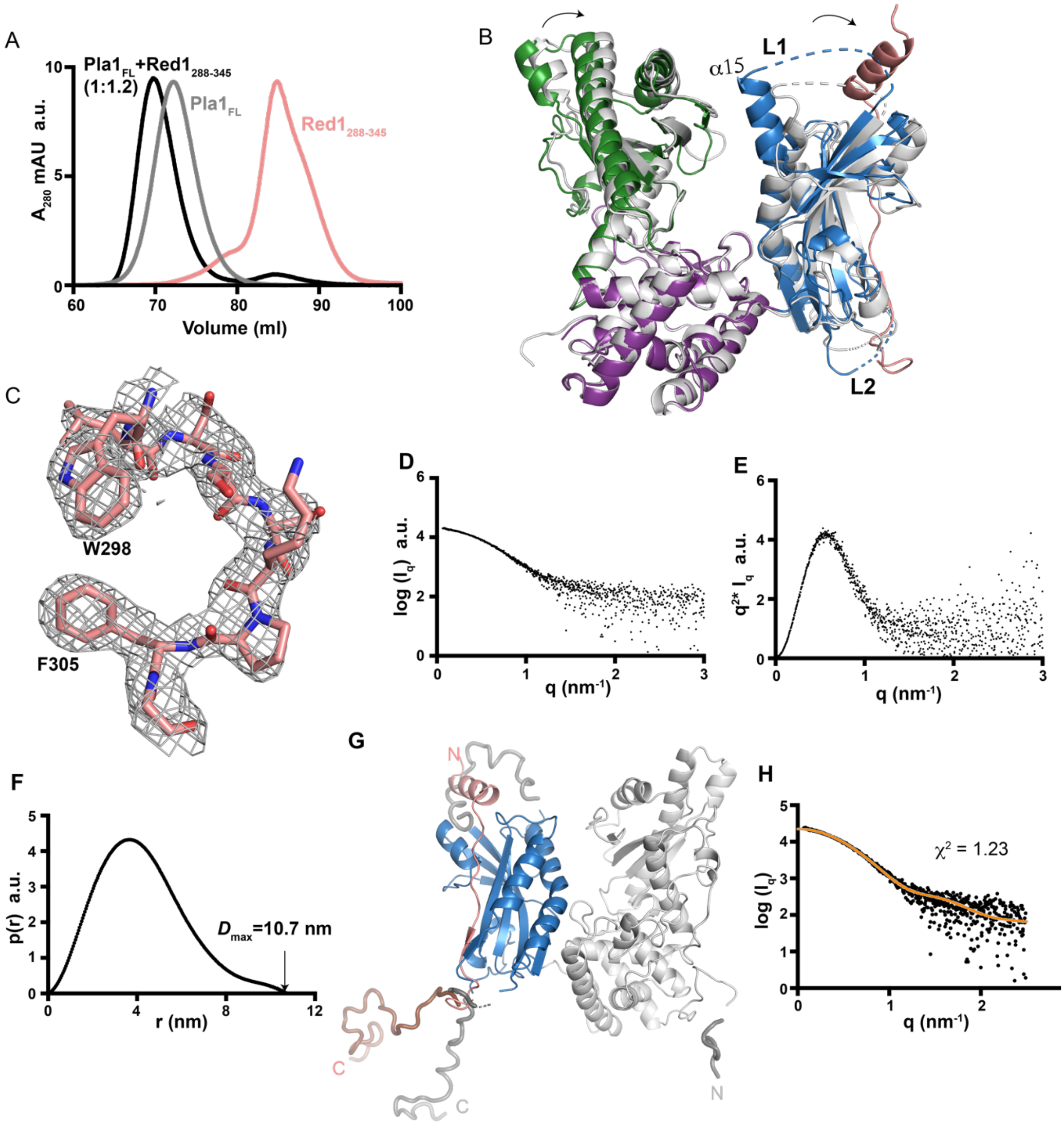


**Supplementary Figure S5. Assembly Pla1-Red1 complex**

(A) Size-exclusion chromatography profiles of Red1$_{288\text{-}345}$ (salmon), Pla1$_{FL}$ (grey) and Pla1$_{FL}$+Red1$_{288\text{-}345}$ (molar ratio 1:1.2 of Pla1$_{FL}$: Red1$_{288\text{-}345}$, shown in black). A marked change in the retention volume of the Pla1$_{FL}$+Red1$_{288\text{-}345}$ complex compared to the free Pla1$_{FL}$ and Red1$_{288\text{-}345}$ is observed. a.u. represent arbitrary units. (B) Structure superposition of Pla1$_{FL}$ and Pla1$_{FL}$+Red1$_{288\text{-}345}$ complex based on alignment of the MD shows a clockwise displacement of NTD and RRM domains as marked by arrows. (C) 2Fo-Fc map of Pla1-Red1 complex with Red1 residues

Trp298-Phe305 contoured at 1σ shown. (D) SAXS profile plotted as log($I_q$) versus q (E) kratky plot and (F) pairwise distribution p(r) curve of $Pla1_{FL}$+$Red1_{288-345}$ complex are presented. a.u. represents arbitrary units. (G) Structure model of $Pla1_{FL}$+$Red1_{288-345}$ complex generated using CORAL[1] to model residues (represented as tubes) disordered in the crystal structure. The NTD and MD are shown in grey while the RRM domain and Red1 are shown in blue and salmon, respectively. (H) Fitting between experimental SAXS and back-calculated SAXS profiles of the $Pla1_{FL}$+$Red1_{288-345}$ complex model generated from CORAL as calculated using CRYSOL[2] and the $\chi^2$ fitting-value of 1.23 are reported.

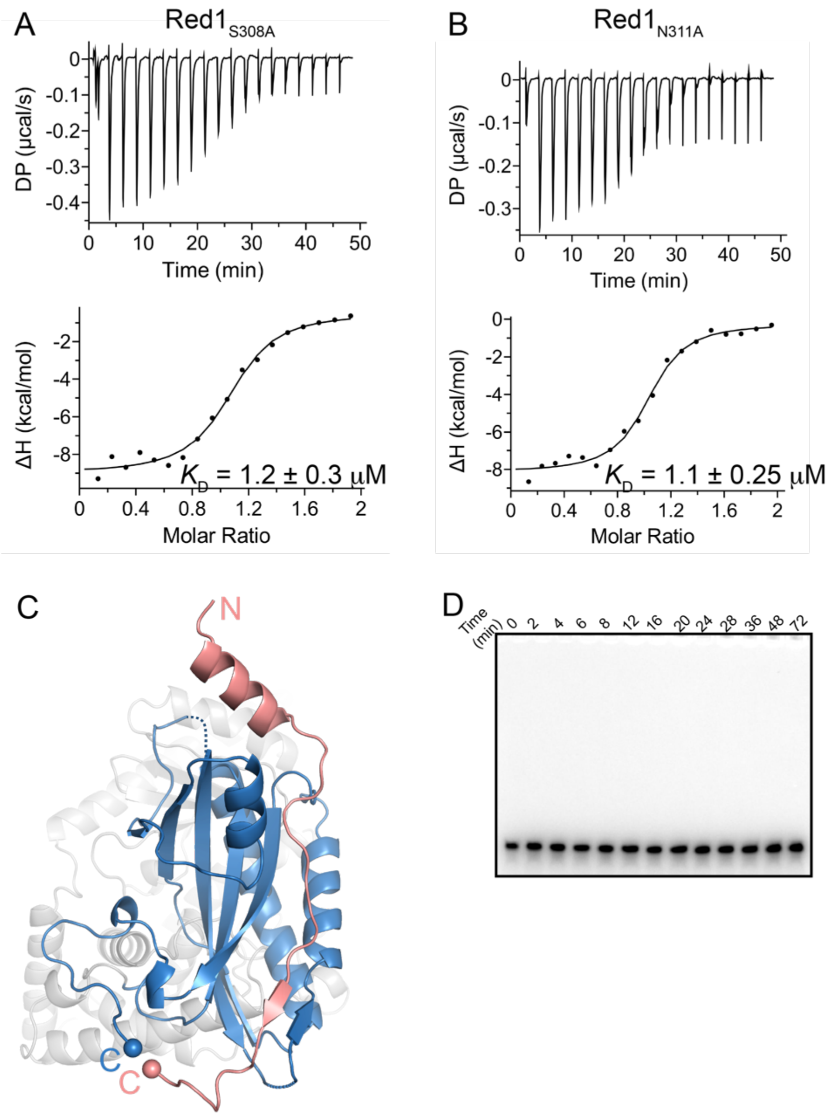


**Supplementary Figure S6. Mutational analysis of Pla1-Red1 complex**

(A) Isothermal titration calorimetry experiments with specific point mutants of Red1$_{288\text{-}345}$ (S308A, N311A) serially titrated into Pla1$_{RRM}$ domain are presented. The calculated dissociation constant ($K_D$) from an average of two independent measurements is shown. (B) Spatial proximity between the C$_\alpha$ atoms of Pla1 residue Ala542 and Red1 residue Gly322 which are the last residues at the C-termini of the proteins visible in the crystal structure of the complex are shown as spheres and the distance between them is 3.5 Å. (D) Polyadenylation of a 5'-Cy3 labeled A$_{15}$ RNA primer by a catalytic mutant of Pla1 (D153A) analyzed by 14% denaturing urea PAGE at different time points.

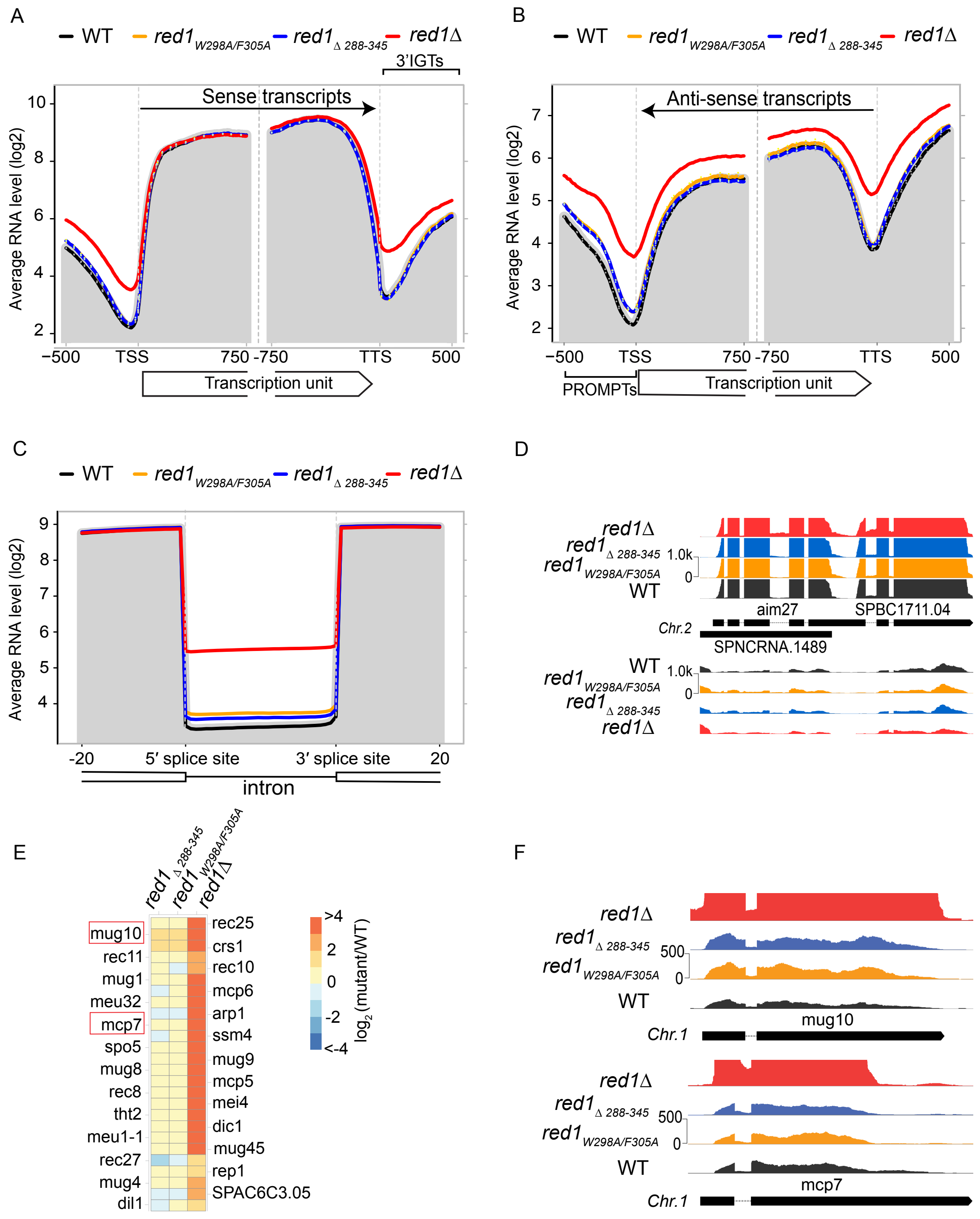


**Supplementary Figure S7. Pla1 in the context of the MTREC complex is responsible for degradation of PROMPTs.**

(A and B) Metagene profile of sense (A) and antisense (B) RNA levels in the indicated strains for all of *S. pombe* genes. The plots represent the geometric average RNA levels from 500bp upstream to 750bp downstream of the transcription start site (TSS) and 750bp upstream to 500bp downstream of the transcription termination site (TTS). Solid lines represent the average of two replicates for all indicated strains, with the exception of *red1*Δ which represent a single dataset. Dotted lines represent the individual biological replicates. The grey shading represents the average RNA levels in the WT strain. (C) Metagene profile of sense RNA levels in the indicated strains for all of *S. pombe* introns. The plots represent the geometric average of RNA values from 20bp upstream to 20bp downstream of annotated introns, with the intronic regions scaled to the same lengths for all introns (stretched or condensed). The grey shading represents the average RNA levels in the WT strain. (D) Strand-specific RNA-seq read coverage of a representative set of genes containing multiple introns, in WT and mutant strains. (E) Heat map representation of changes in RNA levels for the meiotic gene cluster in indicated mutant strains compared to WT ($log_2$ scale). (F) Strand-specific RNA-seq read coverage of a representative set of meiotic genes (Mug10, Mcp7) in WT and mutant strains.

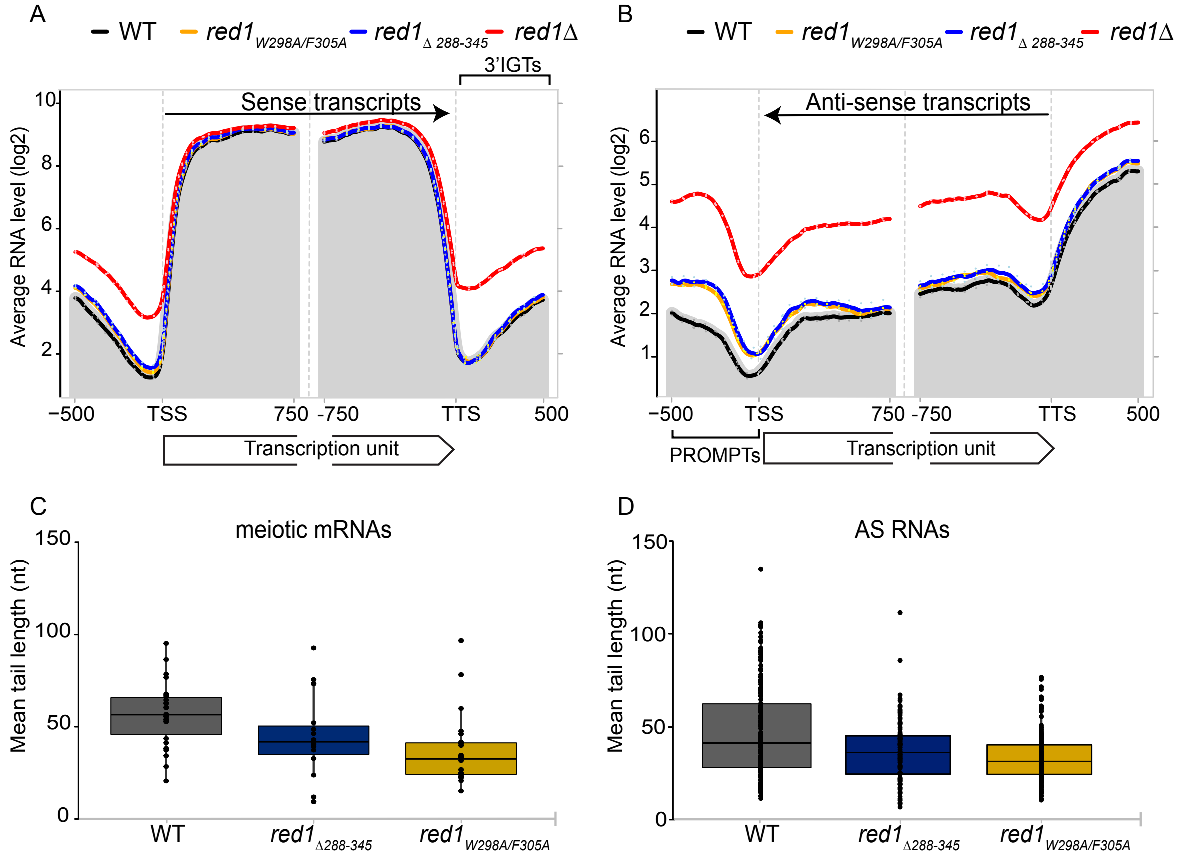


**Supplementary Figure S8. Pla1 in the context of the MTREC complex is responsible for extending the poly(A) tail of CUTs.**

(A and B) Metagene profile of sense (A) and antisense (B) RNA levels in the indicated strains for a subset of *S. pombe* genes with detectable levels of PROMPTs (2400 genes). The strand-specific total RNA seq experiments were carried out without poly(A) selection, using random hexamers for reverse transcription (see details in Methods section). The plots represent the geometric average RNA levels from 500bp upstream to 750bp downstream of the transcription start site (TSS) and 750bp upstream to 500bp downstream of the transcription termination site (TTS). Solid lines represent the average of two replicates for all indicated strains. Dotted lines represent the individual biological replicates. The grey shading represents the average RNA levels in the WT strain. (C and D) Box-plot of mean poly(A) tail length distribution of meiotic mRNAs (C) and AS RNAs (D) for the WT and indicated mutant strains. Dots represent the mean poly(A) tail length of individual meiotic mRNAs/AS RNAs, boxes represent the 25-75 percentile range and the lines represent the median values.

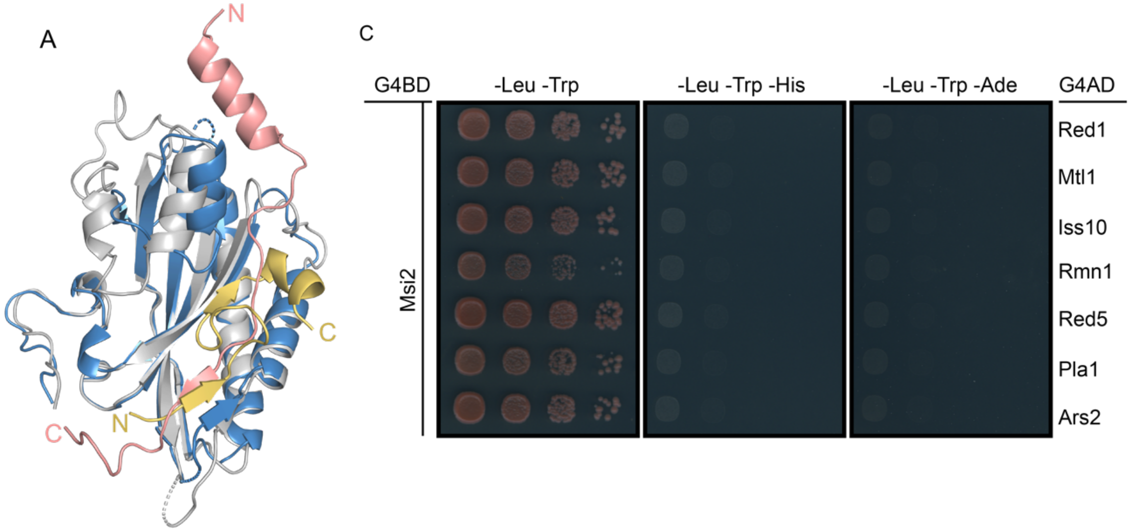


B

```
Fip1    1   ....MSSSEDEDDKFLYGSDSELALPSSKRSRDDEADAGASSNPDIVKRQKFDSPVEETP
Iss1    1   MSNADQHMDVDEDEYLYGETIEERN........DGVT.......................
            58                                                      110
Fip1   57   ATARDDRSDEDIYSDSSDDDSDSDLEVIISLGPDPTRLDAKLLDSYSTAATSSSKDVISV
Iss1   30   ..VSNA....KSPEQASEESDDSDIEFIIETKPGERAE..PLGGSIATLGST........

Fip1  117   ATDVSNTITKTSDERLITEGEANQGVTATTVKATESDGNVPKAMTGSIDLDKEGIFDSVG
Iss1   74   ..............RPS..AKPQVEKTAVEVKTTEPQDLSTAETAPKVDIDAVPTIDGKN

Fip1  177   ITTIDPEVLKEKPWRQPGANLSDYFNYGFNEFTWMEYLHRQEKLQQDYNPRRILMGLLSL
Iss1  118   IFEIDLESFDDKPWRKPGADISDYFNYGFDEFTWAAYCAKQTTLRDDFSPQKFMASLAQI

Fip1  237   QQQGKLNSANDTDSNLGNIIDNNNNVNNANMSNLNSNMGNSMSGTPNPPAPPMHPSFPPL
Iss1  178   PG..PLPSGNVPSP..............A...EFQAMMA...SG.....FPPFMPIPPPI

Fip1  297   ....PMFGSFPPFPMPGMMP............PMNQQPNQNQNQNSK.............
Iss1  211   GGPQEDMGYSRNFPVHASSNYNTTHTSGGGVHSGAATPNAYVNNNPSSSRRESESPANSP

Fip1        ............................................................
Iss1  271   NITSSAGMTHAQPTHNPTSSYGNGASTNYNASRPPSNHPHSSNYPSSSRRKPSPDRYSNY

Fip1        ..............
Iss1  331   SSRGSGGRYRRNRY
```

**Supplementary Figure S9. Interaction between MTREC and CPF**

(A) Structure superposition of *S. cerevisiae* Pap1 (grey) - Fip1 (dark yellow) complex (PDB ID: 3C66) and Pla1 (blue) - Red1 (salmon) complex. Only the RRM domains of Pap1 and Pla1 are shown. (B) Sequence alignment between Fip1 and Iss1. Residues which are identical are highlighted in red while those that are highly similar are colored in red and boxed in blue. Fip1 residues 58-110 shown to be important for binding to Pap1 in *S. cerevisiae*[3] are marked and the corresponding Iss1 residues

30-76 are marked in grey. (C) Y2H experiments do not show interaction between Msi2 and MTREC components apart from Mmi1 and Pab2- related to Figure 7F.

**Supplementary Table S1.** SAXS data collection, processing and modeling statistics

| | **Pla1-Red1 complex** |
|---|---|
| **Data collection parameters** | |
| Instrument | PETRA III, P12 DESY |
| Beam geometry ($mm^2$) | 0.2 × 0.12 |
| Wavelength (Å) | 0.124 |
| $q$ range ($nm^{-1}$) | 0.00226 - 7.4054 |
| Exposure time (s) | 7.8s (40×0.195s) |
| Temperature (°C) | 20 |
| Concentration range measured (mg $ml^{-1}$) | 0.05 - 1.7 |
| Concentration used (mg $ml^{-1}$) | 0.85 |
| **Structural parameters** | |
| $R_g$ (nm) (from $P(r)$) | 3.28 |
| $R_g$ (nm) (from Guinier) | 3.3±0.02 |
| $D_{max}$ (nm) | 10.7 |
| Porod volume estimate ($nm^3$) (from P(r)) | 137.79 |
| **Molecular weight determination (kDa)** | |
| From volume of correlation ($V_c$) | 90.7 |
| From Bayesian assessment [credibility interval], probability | 91.175 [86.95, 95.8], 94.6% |
| Calculated *MW* from sequence | 73 |
| **Software employed** | |
| Primary data reduction | SASFLOW |
| Data processing | PRIMUS |
| Flexibility modeling | CORAL |
| Computation of model intensities | CRYSOL |
| 3D graphics representations | PYMOL |

**Supplementary Table S2: BLI kinetics experiments**

| Biotinylated ligand | Analyte | $K_D$ (nM) | $k_{on}$ (1/Ms) | $k_{off}$ (1/s) | $\chi^2$ | $R^2$ |
|---|---|---|---|---|---|---|
| $Red1_{288-345}$ | $Pla1_{FL}$ | 59.2±0.57 | $2.75*10^5 \pm 2.6*10^3$ | $1.63*10^{-2} \pm 2.9*10^{-5}$ | 2.43 | 0.9949 |
| Iss1 | $Pla1_{FL}$ | 115.2±1.6 | $2.1*10^5 \pm 2.8*10^3$ | $2.42*10^{-2} \pm 5.8*10^{-5}$ | 1.84 | 0.9946 |

**Supplementary Table S3: *E. coli* expression, yeast two hybrid plasmids and yeast strains used in this study**

***E. coli* expression plasmids**

| Plasmid No. | Name | Selection | Reference |
|---|---|---|---|
| ND766 | pET_His-spPla1-1-566 | Kan | this study |
| ND788 | pET_His-spPla1-1-542 | Kan | this study |
| ND1119 | pET_His-1a-spPla1-352-566 | Kan | this study |
| ND940 | pET_Gb1_CHis-spRed1-240-345 | Amp | this study |
| ND953 | pET_Gb1_CHis-spRed1-288-345 | Amp | this study |
| ND954 | pET_Gb1_CHis-spRed1-288-322 | Amp | this study |
| ND82 | pET_MBP_1a-EYFP | Kan | Gunter Stier |
| ND769 | pET21d_spPla1-352-566 | Amp | this study |
| ND775 | pET_MBP_1a- spRed1-240-345 | Kan | this study |
| ND809 | pET_MBP_1a- spRed1-259-288 | Kan | this study |
| ND810 | pET_MBP_1a- spRed1-288-322 | Kan | this study |
| ND811 | pET_MBP_1a- spRed1-288-345 | Kan | this study |
| Rp01 | pET_Gb1_CHis-spRed1-RSRR | Amp | this study |
| Rp02 | pET_Gb1_CHis-spRed1-S308A | Amp | this study |
| Rp04 | pET_Gb1_CHis-spRed1-WFAA | Amp | this study |
| Rp05 | pET_Gb1_CHis-spRed1-N311A | Amp | this study |
| Rp10 | pET_His-1a-spPla1-EEQE | Kan | this study |
| Rp11 | pET_His-1a-spPla1-EEE | Kan | this study |
| Rp12 | pET_His-spPla1-D153A | Kan | this study |
| Rp17 | pET_Gb1_CHis-spRed1-V313E | Amp | this study |

| | | | |
|---|---|---|---|
| Rp18 | pET_His-1a-spPla1-K368E | Kan | this study |

**Yeast two hybrid plasmids**

| Plasmid No. | Name | Selection | Reference |
|---|---|---|---|
| ND333 | pGBKT7 | Kan | Dobrev et al.[4] |
| ND334 | pGADT7 | Amp | Dobrev et al.[4] |
| ND368 | pGADT7-spRed1_1-712 | Amp | Dobrev et al.[4] |
| ND370 | pGADT7-spMtl1_1-1030 | Amp | Dobrev et al.[4] |
| ND335 | pGADT7 -spRed5_1-376 | Amp | Dobrev et al.[4] |
| ND336 | pGADT7 -spIss10_1 -551 | Amp | Dobrev et al.[4] |
| ND337 | pGADT7 -spMmi1_1-488 | Amp | Dobrev et al.[4] |
| ND338 | pGADT7 -spPab2_1 -166 | Amp | Dobrev et al.[4] |
| ND339 | pGADT7 -spRmn1_1 -590 | Amp | Dobrev et al.[4] |
| Rp19 | pGADT7 -spPla1_1 -566 | Amp | this study |
| Rp20 | pGADT7 -spArs2_1-609 | Amp | this study |
| Rp21 | pGBKT7 -spMsi2_1 -474 | Kan | this study |
| Rp15 | pGADT7-spRed1_1-712-WFAA | Amp | this study |
| Rp16 | pGADT7-spRed1_del288-345 | Amp | this study |

**Yeast strains**

| Strain No. | Name | Relevant Genotype | Reference |
|---|---|---|---|
| P419 | WT | *MatMsmt0, leu1-32, ura4-D18, ade6-216* | |
| P1 | WT | *MatMsmt0, leu1-32, ura4 DS/E, ade6-210, his2, Otr1R:: Ura4 (SphI)* | |
| P344 | WT | *h+, HIS, leu1-32, ade6-210, ura4-D18* | V2-33-H12, Bioneer Inc |
| F3230 | Red1 (WT) | *MatMsmt0, leu1-32, ura4-D18* | this study |

| | | | |
|---|---|---|---|
| | | *ade6-M210, Red1:: natNT2 (+3'UTR)* | |
| F3232 | *red1*$_{W298A/F305A}$ | *MatMsmt0, leu1-32, ura4-D18 ade6-M210/M216, Red1*$_{W298A/F305A}$ *:: natNT2 (+3'UTR)* | this study |
| F3436 | *red1*$_{\Delta 288\text{-}345}$ | *MatMsmt0, leu1-32, ura4-D18, ade6-M210, Red1*$_{\Delta 288\text{-}345}$ *:: natNT2 (+3'UTR)* | this study |
| F3237 | *red1Δ* | *MatMsmt0, leu1-32, ura4-D18, ade6-M210, red1Δ:: kanMX6* | this study |
| F3269 | Red1-FTP | *MatMsmt0, leu1-32, ura4-D18, ade6-M216, Red1-3xFTP:: hyg* | this study |
| F3271 | *red1*$_{W298A/F305A}$*-FTP* | *MatMsmt0, leu1-32, ura4-D18, ade6-M210, Red1*$_{W298A/F305A}$*-3xFTP::hyg* | this study |
| F3273 | *red1*$_{\Delta 288\text{-}345}$*-FTP* | *MatMsmt0, leu1-32, ura4-D18, ade6-M216, Red1*$_{\Delta 288\text{-}345}$*-3xFTP::hyg* | this study |
| F3043 | Red1-FTP | *h90, leu1-32, ura4-D18, ade6-M210, Red1- 3xFTP::hyg.* | this study |
| F3197 | *red1*$_{W298A/F305A}$*-FTP* | *h90, leu1-32, ura4-D18, ade6-M210 Red1*$_{W298A/F305A}$*-3xFTP::hyg.* | this study |
| F3199 | *red1*$_{\Delta 288\text{-}345}$*-FTP* | *h90, leu1-32, ura4-D18 ade6-M210, Red1*$_{\Delta 288\text{-}345}$*-3xFTP::hyg.* | this study |
| F3203 | Pla1-HA | *h+, HIS, leu1-32, ade6-210, ura4-D18 Pla1-3xHA::kanMX6* | this study |

| | | | |
|---|---|---|---|
| F3205 | Red1-FTP; Pla1-HA | *h90, leu1-32, ura4-D18, ade6-M210 Red1-3xFTP::hyg; Pla1-3xHA::kanMX6* | this study |
| F3207 | *red1$_{W298A/F305A}$-FTP*; Pla1-HA | *h90, leu1-32, ura4-D18, ade6-M210 Red1$_{W298A/F305A}$-3xFTP::hyg; Pla1- 3xHA::kanMX6* | this study |
| F3209 | *red1$_{\Delta 288\text{-}345}$-FTP*; Pla1-HA | *h90, leu1-32, ura4-D18, ade6-M210 Red1$_{\Delta 288\text{-}345}$-3xFTP::hyg; Pla1-3xHA::kanMX6* | this study |

### Supplementary Table S4: Primers used in this study

| Primer No. | *Name* | Sequence (5’->3’) |
|---|---|---|
| ND342 | *spPla1-NcoI-FW* | TATACCATGGGTACTACCAAGCAATGGGGTATTACAC |
| ND343 | *spPla1-BamHI-RE* | TATAGGATCCTTATGCCGTTGAAACTTTTTGTCGTTTTAATTG |
| ND432 | *spPla1-352-NcoI-FW* | TATACCATGGACTTTTTTCATCGTTATAAGCATTATC |
| ND433 | *spPla1-352-XhoI-RE* | TATACTCGAGTTAGTCGTGTTTTTGAAACAAAGCTGAC |
| N748 | *spPla1-542-STOP-BamHI-RE* | TATAGGATCCTTAAGCTTTCGGCCGTTCTTCTCC |
| N620 | *spRed1_288-FW-GA-BamHI* | GAAGAACGGCCGAAAGCTGGTGGTTCTGGCGGTTCCATATCTTTACCACTTTTGAAGCAGG |
| N619 | *SpRed1_345-RE-GA-BamHI* | GTCAGTGGTGGTGGTGGTGGTGGGATCCATTTTCACTATTTGAGGGGG |
| N621 | *SpRed1_322-RE-GA-BamHI* | GTCAGTGGTGGTGGTGGTGGTGGGATCCATCATCATCGGAATCAAATTCAATAAC |

| | | |
|---|---|---|
| KS1 | *spRed1_F305A_FP* | CGATTGGCTATCTTCTTCGAAGCCTGCTGGCTCCTCGAC |
| KS2 | *spRed1_F305A_RP* | AGTCGAGGAGCCAGCAGGCTTCGAAGAAGATAGCCAATCG |
| KS3 | *spRed1_W298A_FP* | CCACTTTTGAAGCAGGACGATGCGCTATCTTCTTCGAAGC |
| KS4 | *spRed1_W298A_RP* | CTTCGAAGAAGATAGCGCATCGTCCTGCTTCAAAAGTGG |
| KS5 | *spRed1_N311A_FP* | GCCTTTTGGCTCCTCGACTCCAGCTGTAGTTATTGAATTTGATTC |
| KS6 | *spRed1_N311A_RP* | GAATCAAATTCAATAACTACAGCTGGAGTCGAGGAGCCAAAAGGC |
| KS7 | *spRed1_D317RRR_FP* | CTTTTGGCTCCTCGACTCCAAATGTAGTTATTGAATTTCGTTCCCGTCGTGATGGAGATGACTTTTCGA |
| KS8 | *spRed1_D317RRR_RP* | TCGAAAAGTCATCTCCATCACGACGGGAACGAAATTCAATAACTACATTTGGAGTCGAGGAGCCAAAAG |
| KS9 | *spRed1_S308A_FP* | CGAAGCCTTTTGGCTCCGCGACTCCAAATGTAGTT |
| KS10 | *spRed1_S308A_RP* | AACTACATTTGGAGTCGCGGAGCCAAAAGGCTTCG |
| KS11 | *spPla1-K368E-RP* | GAGCTTCAGCTGTCTCAGCAGCTGCTGTGATCG |
| KS12 | *spPla1-K368E-FP* | CGATCACAGCAGCTGCTGAGACAGCTGAAGCTC |
| KS13 | *spPla1_EEE_FP* | AAGAACGGCCGAAAGCTACTGAGGAGGAGTCCACTGCTGACACTGCAC |
| KS14 | *spPla1_EEE_RP* | GTGCAGTGTCAGCAGTGGACTCCTCCTCAGTAGCTTTCGGCCGTTCTT |
| KS15 | *spPla1_EEQE_BamHI_RP* | CATGTCGGATCC TTA TGCCGTTGAAACTTCTTGTTCTTCTAATTGCTCTGTACTATG |
| KS16 | *spPla1_352_NcoI_FP* | GAGACTACC ATGG ACTTTTTTCATCGTTATAAGCATTATCTTAC |
| KS17 | *spPla1_D153A_FP* | AGTTTAAATTTTTGGGAATAAGTATAGCTTTAATTTTTGCTCGTCTTTCAGTTCC |

| | | |
|---|---|---|
| KS18 | *spPla1_D153A_RP* | GGAACTGAAAGACGAGCAAAAATTAAAGCTATACTTATTCCCAAAAATTTAAACT |
| KS19 | *spRed1-V313R-RP* | CATCGGAATCAAATTCAATACGTACATTTGGAGTCGAGGAGCC |
| KS20 | *spRed1-V313R-FP* | GGCTCCTCGACTCCAAATGTACGTATTGAATTTGATTCCGATGA |
| KS21 | *spIss1-30-NcoI-FP* | GAGACTACC ATG G TGTCAAATGCCA AGTCACCAGA |
| KS22 | *spIss1-76-BamHI-RP* | TATAGGATCCTTAAGAAGGTCGGGTTGAACCTAATGTT |
| KS23 | *spRed1-del288-345-RP* | GTAATCTGAACGAGACATAGTAAGAAGATTTTTTCTTTCAGGAAGGACG |
| KS24 | *spRed1-del288-345-FP* | CGTCCTTCCTGAAAGAAAAAATCTTCTTACTATGTCTCGTTCAGATTAC |
| KS25 | *spMsi2_NcoI_FW-GBKT7--GA* | AGGAGGACCTGCATATGGCCATGGGCGGCTCTGATTTTGAAGATG |
| KS26 | *spMsi2_BamHI_RE-GBKT7-GA* | TGCAGGTCGACGGATCCTTATCGACGATATGGATGGAAGCTGTGCC |
| KS27 | *spPla1_NcoI_FW-ADT7-GA* | GGCGAGCGCCGCCATGACTACCAAGCAATGGGGTATTACAC |
| KS28 | *spPla1_BamHI_RE-ADT7-GA* | GCAGCTCGAGCTCGATGGATCCTTATGCCGTTGAAACTTTTTGTCGTTTTAATTG |
| KS29 | *spArs2_NcoI_FW-ADT7-GA* | GGCGAGCGCCGCCATGGCTAGTGAGGTCCATCAAGAAAG |
| KS30 | *spArs2_BamHI_RE-ADT7-GA* | GCAGCTCGAGCTCGATGGATCCTTAGTAATCTAATTCTGGAACCTCCTGGTTAG |
| AN77 | *spRed1_For_ F305A* | GCGCTATCTTCTTCGAAGCCTGCTGGCTCCTCGACTCCAAATG |

| | | |
|---|---|---|
| AN78 | *spRed1_Rev_W298A* | GCAGGCTTCGAAGAAGATAGCGCATCGTCCTGCTTCAAAAGTGG |
| AN80 | *spRed1_ For_Nhe1 site* | CGGACTGGATTCCTCTTCTGCTAGCTCCCAAGGAAGTCTGAC |
| AN70 | *spRed1-Pla1 DEL_Rev_first frag* | CTGAACGAGACATAGTAAGACCAAGATTTTTTCTTTCAGGAAGG |
| AN81 | *spRed1_Rev with Kpn1 site* | GATATAATCGACAAGCGGTACCTTATCCTCAAAATCTGTTTCCGTC |
| AN82 | *spRed1_For DEL sec_ frag.* | CGTCCTTCCTGAAAGAAAAAATCTTGGTCTTACTATGTCTCGTTCAG |
| AN62 | *spRed1_Fw_above 400bp* | CGCTTGTCGATTATATCTCTC |
| AN63 | *spRed1_Rev_natNT2OH* | CGTCGACCTGCAGCGTACGTTATCAATAACCATTGAAATG |
| AN64 | *spRed1_Fw_ Clonat Overhang* | CGAGCTCGAATTCATCGATAAATGCGCGTATTGTAATAAG |
| AN65 | *spRed1_ Rev. 500bp after.* | GTTTGGAAGATGGTTTACGCCTAC |
| AN86 | *Pla1-3xHA tag first frag._For.* | GCTCAACAAGTAGCTAGCGG |
| AN87 | *Pla1-3xHA tag first frag._Rev.* | GTTAATTAACCCGGGGATCCGTGCCGTTGAAACTTTTTGTC |
| AN88 | *Pla1-3xHA tag sec. frag._For.* | CGAGCTCGAATTCATCGATGTAAATACCCACACATAAGACTC |
| AN89 | *Pla1-3xHA tag sec. frag._Rev.* | GAACACTCCTAAAGATCCCAAC |
| AN139 | *ChIP_check_ura4DS/E#1* | GAGGGGATGAAAAATCCCAT |
| AN140 | *ChIP_check_ura4DS/E#2* | TTCGACAACAGGATTACGACC |
| AN162 | *GAPDH_For.* | AACATCATCCCCTCCTCCAC |
| AN163 | *GAPDH_Rev.* | GCCTTGATGTCCTCGTAGTTG |

### Supplementary references:


1. Petoukhov, M.V. et al. New developments in the ATSAS program package for small-angle scattering data analysis. *J Appl Crystallogr* **45**, 342-350 (2012).

2. Franke, D. et al. ATSAS 2.8: a comprehensive data analysis suite for small-angle scattering from macromolecular solutions. *J Appl Crystallogr* **50**, 1212-1225 (2017).
3. Kumar, A. et al. Dynamics in Fip1 regulate eukaryotic mRNA 3' end processing. *Genes Dev* (2021).
4. Dobrev, N. et al. The zinc-finger protein Red1 orchestrates MTREC submodules and binds the Mtl1 helicase arch domain. *Nat Commun* **12**, 3456 (2021).
